# Emergence of persistent and sequential representations in the hippocampal-prefrontal circuitry during associative learning

**DOI:** 10.1101/2022.06.28.497926

**Authors:** Bryan C. Souza, Jan L. Klee, Luca Mazzucato, Francesco P. Battaglia

**Author notes:** Correspondence (B.C.S.), (L.M.), (F.P.B.). These authors contributed equally to this work (Luca Mazzucato, Francesco P. Battaglia).

## Abstract

Temporal associations between sensory stimuli separated in time rely on the interaction between the hippocampus and medial prefrontal cortex (mPFC). However, it is not known how changes in their neural activity support the emergence of temporal association learning. Here, we use simultaneous electrophysiological recordings in the hippocampal CA1 region and mPFC of mice to elucidate the neural dynamics underlying memory formation in an auditory trace conditioning task. We found that in both areas conditioned (CS+/CS−) and unconditioned stimuli (US) evoked similar temporal sequences of neural responses that progressively diverged during learning. Additionally, persistent CS representations emerged in mPFC after learning, supported by CS+ coding states whose transient reactivation reliably predicted lick onset and behavioral performance on single trials. These results show that coordination of temporal sequences in CA1 and persistent activity in mPFC may underlie temporal association learning, and that transient reactivations of engrams in mPFC predict the animal behavior.

**Highlights:**

- Temporal representation of CS+, but not CS− and US, strengthens in CA1 after learning.
- Similarity between stimulus and reward temporal representations decrease with learning, but is recovered in error trials.
- Representation of stimulus identity is strong and stable in PFC since stimulus onset, while it only emerges in CA1 during trace period.
- Neural states defined on faster time-scales reveal the emergence of CS coding states in PFC whose onset predicts lick times and task performance.
- PFC-CA1 states do not increase coordination during late CS+ stimulus and trace.

Figure 0:
Graphical Abstract

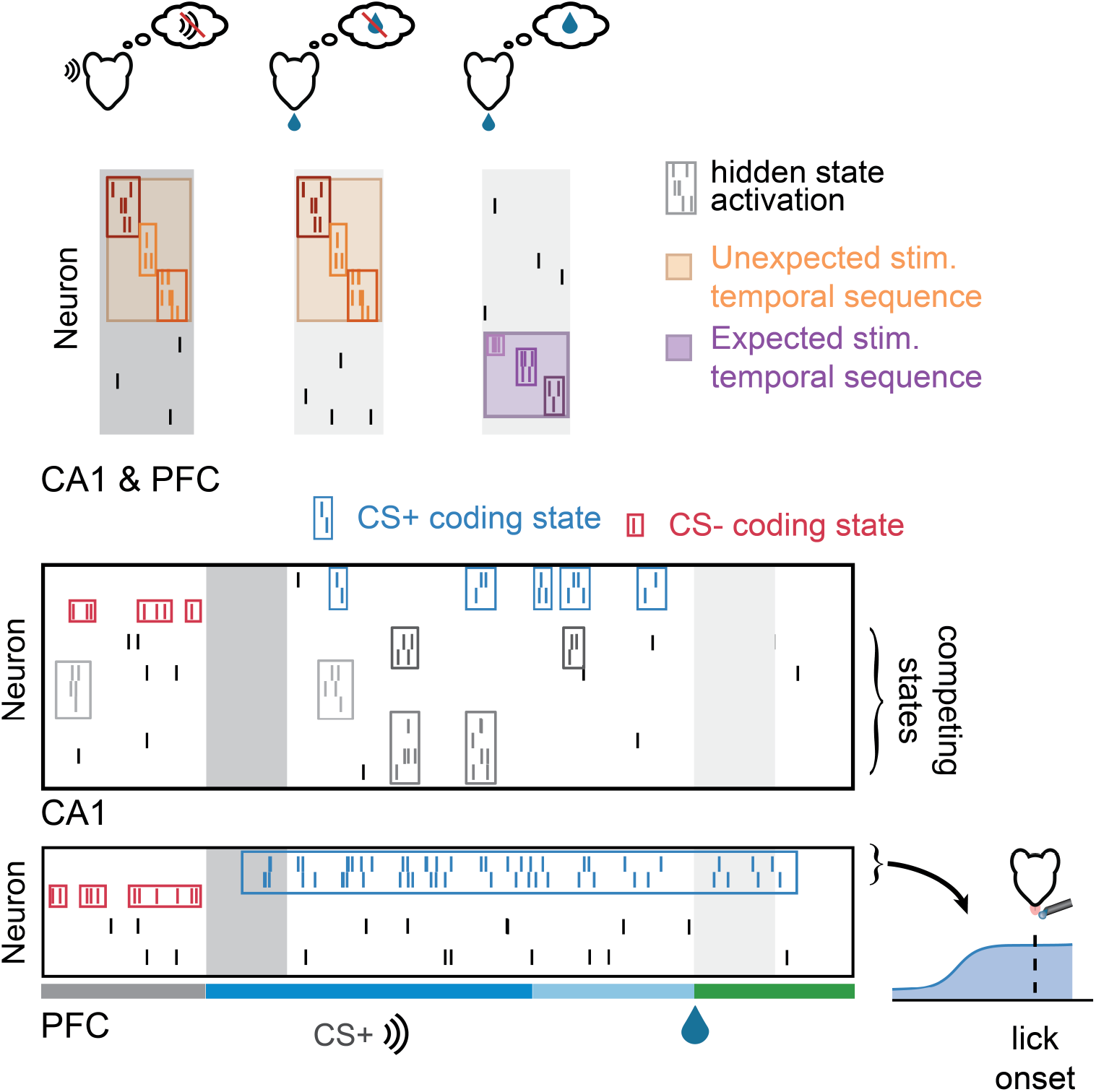

## Introduction

The hippocampus and the medial prefrontal cortex (mPFC) are both known to have important roles in sensory and memory representations. For instance, the hippocampus integrates multiple sensory inputs coming from the entorhinal cortex, bridging the ’what’, ’where’ and ’when’ components of a memory (Eichenbaum (2017a)) while the mPFC is believed to create the stimulus-context associations that support decision making (Euston et al. (2012)) and to support working memory via sustained attention on relevant sensory representations (Postle (2006), Lara and Wallis (2015)). Although the interaction between those two areas are known to support different cognitive processes (Battaglia et al. (2011), Eichenbaum (2017b), Shin and Jadhav (2016), Preston and Eichenbaum (2013)), it is still unknown how the interplay between the mPFC and hippocampus can lead to the formation of internal representations linking events separated in time.

The mPFC neuronal activity can be modulated by different stimuli and reward (Starkweather et al. (2018), Otis et al. (2017)). Beyond responding to positive outcomes (Euston et al. (2012), Burton et al. (2009), Gruber et al. (2010), Pratt and Mizumori (2001)), the mPFC can also encode expectancy of both positive and negative outcomes (Baeg (2001), Gilmartin and McEchron (2005)), and reward absence (Quilodran et al. (2008)). Moreover, mPFC is implicated in acquisition of context-dependent rules (Miller and Cohen (2001), Miller (2000)) and required for the maintenance of stimulus representations during delays in working memory tasks (Goldman-Rakic (1995), Funahashi et al. (1993)), even though this representation might be stored elsewhere (Lara and Wallis (2015), Postle (2006)). The mPFC was shown to encode long memory traces of past decisions (Murakami et al. (2017)) and rewards (Bernacchia et al. (2011)) and it is required to implement trial-history biases in decision-making tasks (Murakami et al. (2017).

Similar to the mPFC, the hippocampus is also required in tasks involving delayed decision (Hattori et al. (2015), McEchron and Disterhoft (1999, 1997)). Beyond its strong link to spatial processing (Moser et al. (2015, 2008)), hippocampal activity also encodes multiple temporal features, possibly bridging task events separated in time through temporal sequences (Pastalkova et al. (2008a), MacDonald et al. (2011), Naya and Suzuki (2011), Eichenbaum (2014)). For instance, CA1 inactivation during a trace fear condition task impaired learning by preventing the temporal binding of the CS and US stimuli in memory (Sellami et al. (2017)). During trace fear conditioning, a reorganization of CA1 ensemble activity occurred during learning of the conditioned response, linked to the emergence of sparse CS-selective responses in pyramidal cells (Ahmed et al. (2020)). However, the circuit mechanisms underlying CA1 network reorganization and the role of the CA1-mPFC coordinated interaction during learning are unknown.

In this work, we aimed at elucidating the neural mechanisms leading to the emergence of a temporal association, linking events occurring at different times. To this end, we recorded neurons from the hippocampal area CA1 and mPFC of mice engaged in an appetitive auditory trace-conditioning task (Klee et al. (2021)), and examined how learning shaped the relationship between stimulus responses, temporal encoding, and behavior. We found that both conditioned (CS) and unconditioned stimuli (US) triggered the onset of temporal sequences in both CA1 and mPFC, which in turn were differentially modulated by learning. Moreover, we found, after learning, that unexpected reward delivery evoked an echo of the CS+ temporal sequence in mPFC during the US. We uncovered strong and persistent stimulus representation in the mPFC, which started after the stimulus onset and persisted until the US, bridging stimulus and reward over the trace periods. Using single-trial decoding based on hidden Markov models, we found that the reactivation of CS+ coding states in mPFC predicted the onset of wrong licks in response to CS-stimuli, leading to error trials. Coordination between CA1 and PFC states decreased during CS+ trials, suggesting that CA1 may support internal memory representation during the initial stimulus sampling, while mPFC activity is dominated by network states encoding stimulus-reward mapping and driving behavior in overtrained animals.

## Results

We previously developed an appetitive trace conditioning task (Klee et al, 2021) which trained mice to differentiate between two sound stimuli (conditioned stimulus: CS+ and CS-;Figure 1A). In each CS+ trial, the 2s stimulus was followed by a 1s delay (trace period), and a reward delivered in a water port. The same trial structure was present in CS− trials, except that there was no reward. In order to investigate the neural dynamics in CA1 and mPFC during the trace conditioning task, we acutely inserted high-density silicon probes in CA1 and/or mPFC and recorded spiking activity during the task, before and after learning. Crucially, the presence of a long trace period in this experimental paradigm rules out a simple Hebbian mechanism to link CS and US neural representations, because the two stimuli occur in different epochs of the trial without any temporal overlap Raybuck and Lattal (2014). Additionally, this paradigm allowed us to separate representations that might emerge during the sensory cue, from the internally generated representations that bridge the stimulus to the behavior during the trace period. Learning criteria was achieved when animals licked in anticipation of US in CS+ trials, but not in CS− trials (lick rates statistically different between CS+ and CS− trials; p<0.05, t-test; Figure 1A-C).

**Figure 1:**
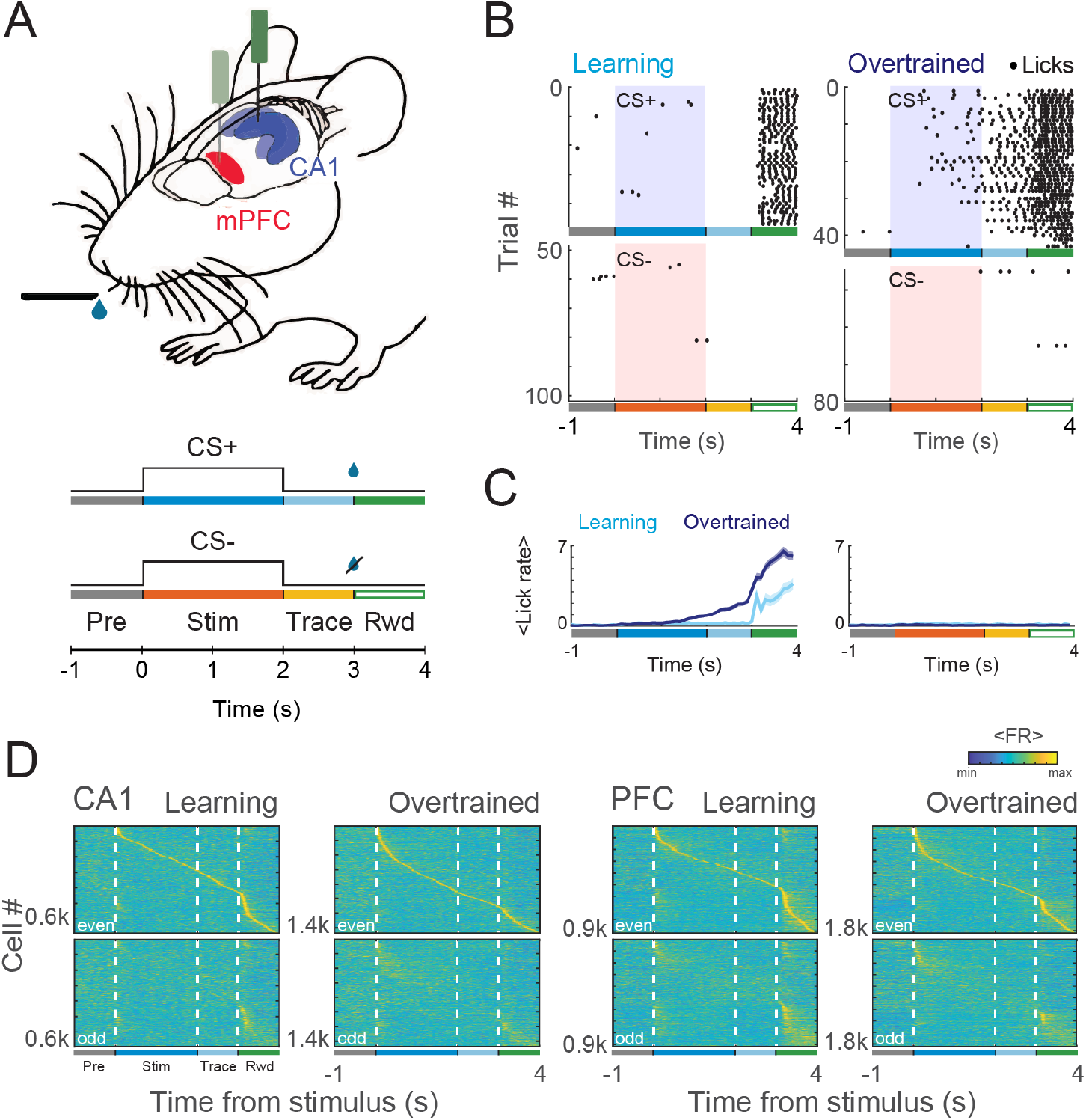
Behavioral and firing rate profile during a trace conditioning. **A.** Schematic representation of the task. Head-fixed animals with acute implants on CA1 and mPFC were presented a 2-sec auditory stimulus (either CS+ or CS-). After a 1-sec (trace period) a reward (US) was available on CS+ trials. **B.** Lick response of an example animal in CS+ and CS− trials before and after learning. **C.** Average lick response for CS+ and CS− trials for all animals. Overtrained sessions were identified by the presence of anticipatory licks in CS+, but not in CS− trials. **D.** Average firing rate profile of all recorded cells over the course of a CS+ trial. Peak firing rate time on even trials was used to sort cells on both groups (even and odd). Note the presence of temporally structured firing after CS and US presentation.

### Differential encoding of temporal information in CA1 and mPFC ensembles

We first investigated whether CA1 and mPFC ensembles encoded the passing of time within the trial by generating sequential activity that consistently tiled the duration of a trial. To test this, we computed the average binned firing rate (80 ms sliding window) of the pooled neurons over a subset of the trials (even trials). The neurons were then sorted according to the time of their peak firing rate. We then applied the same order in the remaining, odd trials to unveil cells firing in a reliable temporal structure. If neurons generated consistent temporal sequences, we would expect the same sequential activation to be present both in the odd and in the even trials. We found a concentration of peak activation around the first 0.5 s after stimulus (CS), which seemed to be modulated by learning (mainly in CA1; Figure 1D). In addition, we also found a concentration of peaks after reward onset (US). Interestingly, some cells peaking in the CS had also a peak at the US and vice-versa (this effect was mainly seen in PFC). Those results unveiled the presence of temporal sequences coordinated by stimulus (CS+) and reward (US) onset at the population level (Figure 1D, Figure S1).

We confirmed these results on a session-by-session basis, using a cross-validated Naive-Bayesian classifier to discriminate time points within a trial from population activity of each session (Figure 2A). Decoding probability was then evaluated for different time bins, building a time-decoding matrix with the corresponding testing bin time in the x-axis and the decoded time on y-axis. If temporal coding was present, we expected a high classification accuracy on the main diagonal where predicted time (y-axis in Figure 2A) matched real time (x-axis). The presence of temporal sequences was then assessed by measuring how much decoding probability successively concentrated in the main diagonal, where decoded time is equal to real sample time.

**Figure 2:**
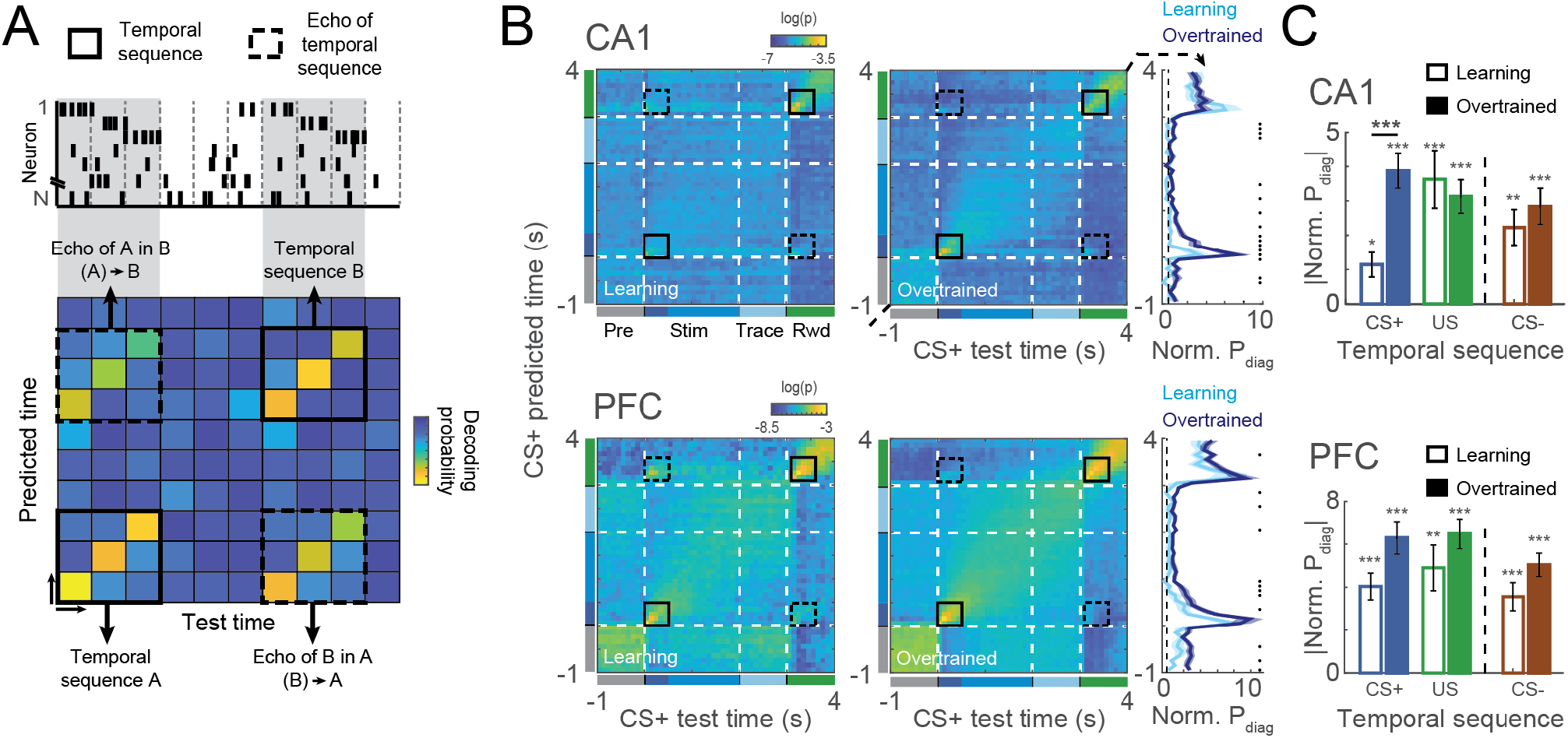
Emergence of temporal sequences for CS+, CS− and US. **A.** Schematic representation temporal sequences in the spike activity (top) and how they emerge in the time decoding matrices (bottom). Bayesian decoders were trained to predict the trial time of a given time bin, which was used to build a matrix with decoding probabilities for each time bin over the course of the trial. Temporal sequences were then identified as clusters of high performance in the main diagonal, where real time matches predicted time. Additionally, diagonal stripes of high temporal decoding performance outside the main diagonal revealed the similarity between two temporal sequences, and were defined as the temporal echo of one sequence (actual time) in the other (predicted time). **B.** Mean time decoding matrices for learning and overtrained sessions together with the average normalized probability on the main diagonal (rightmost panel). Decoders were trained and tested using CS+ activity from CA1 (top) and PFC (bottom). Black boxes show the regions in which temporal sequences (solid lines) and temporal echoes (dashed lines) were present. Black dots denote times in which learning and overtrained decoding are different (Wilcoxon ranksum, p<0.05). **C.** Average (normalized) decoding probability around CS+ and US temporal sequences, taken on main diagonal of the contiguous black boxes in B. Normalization was done by z-scoring probability values using the the mean and s.d. from surrogate decoding matrices. CS− sequences were computed similarly to CS+, but from a decoding matrix computed using CS− activity for training and testing (see also Figure S2). We used aligned rank transform to perform a nonparametric two-way ANOVA using recording area and learning condition on each temporal sequence as factors. We found no significant interaction effect for the two factors (F(1,107)= 0.51, p=0.47; F(1,107)= 0.74, p=0.39 and F(1,107)= 0.64, p=0.43 for CS+, US and CS−, respectively). Simple main effects analysis revealed that PFC had higher temporal sequences than CA1 (F(1,107)= 20.08, p<0.001; F(1,107)= 18.02, p<0.001 and F(1,107)= 14.04, p<0.001 for CS+, US and CS-, respectively). Also, for CS+ temporal sequences, we found an effect on learning condition (F(1,107) = 18.08, p<0.001; F(1,107)= 1.30, p=0.26 and F(1,107)= 3.47, p=0.07 for CS+, US and CS-, respectively). Notice the increase in CS+ temporal representation in CA1 after learning. Shaded areas and errorbars denote SEM. *p<0.05; **p<0.01; ***p<0.001.

Similar to our previous population analysis, we found temporal sequences associated with CS+ and US (Figure 2B). This was evident in the mean time-decoding matrices, but also when looking exclusively to the main diagonal (right-most panel). Interestingly, time-decoding matrices computed using CS− trials also showed a strong temporal sequence associated with the stimulus (Figure S2A). To quantify this effect, we computed the average of the main diagonal in the temporal sequence window (0-0.5s for CS and 3.1-3.6s for US) (Figure 2C). All the temporal sequences (CS+, CS− and US) were significantly different compared to a sham temporal sequence artificially detected before the stimulus onset ((−1)-(−0.5)s).

We then tested whether the strength of temporal encoding differed between areas and whether it was modulated by learning. We found a stronger temporal encoding in mPFC compared to CA1 following each stimulus (non-parametric two-way anova with aligned rank transform (Wobbrock et al. (2011); main effect (mPFC vs. CA1), p<0.001; no interaction effects; see Figure 2C). Moreover, we found stronger CS+ temporal sequence after learning in both areas (main effect (learning vs. overtrained), p<0.001; see Figure 2C). We could also see reduction on the relative dimensionality of the population activity during the temporal sequences in CA1 and (after learning) in PFC (Figure S2B). In general, those results suggest that encoding of time is stronger in PFC than in CA1 and, in particular, during CS+, which is the most task-relevant stimulus. They also imply a link between temporal sequences and learning (Figure 2C; see also Figure S2).

### Temporal sequences of CS+, CS-, US ‘echo’ each other, in a learning-dependent fashion

The time-decoding analysis revealed a surprising ‘echo’ effect, whereby US and CS+ temporal sequences shared a similar representation of time following each stimulus (high classification accuracy in off-diagonal regions, Figure 2A-B). We refer to this effect as the temporal echo of one sequence (the one used to test the model) in another sequence (the predicted sequence).

Because (CS+)–evoked temporal sequences in CA1 were stronger after learning sessions, we hypothesized that temporal echos could be modulated by learning as well. We reasoned that the similarity between US and CS+ representation could be inferred by how well US sequences would generalize to CS+. To investigate this, we computed time-decoding matrices focusing on how US temporal sequences are decoded by a model restricted to the rest of the trial (pre-stimulus, stimulus and trace period; Figure 3A), from CS+ trials.

**Figure 3:**
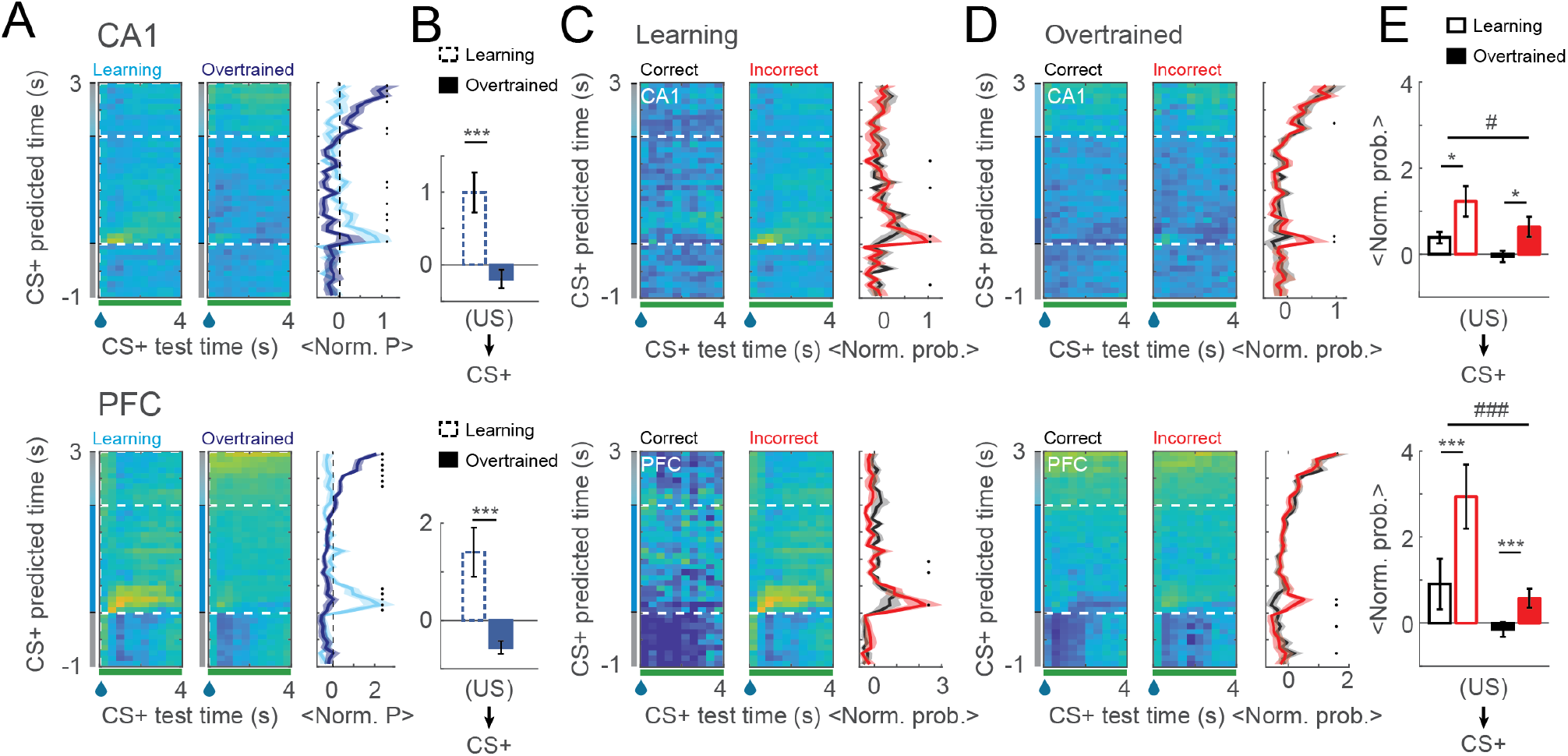
Similarity between CS+ and US temporal representations is modulated by learning and trial outcome. **A.** Time decoding matrices trained using CS+ activity before the US, and used to decode US time (left and middle). Average (normalized) probabilities for the US period are shown for learning and overtrained sessions (right). **B.** Mean decoding in the temporal echo window for CA1 and PFC. **C-D.** Timedecoding matrices as in A, but showing the temporal echo in correct and incorrect trials from learning (C) and overtrained (D) sessions. **E.** Average performance on the temporal echo window (main diagonal) for CA1 (top) and PFC (bottom). We used aligned rank transform to perform a nonparametric two-way ANOVA using trial outcome (i.e., correct vs. incorrect) and learning condition as factors for each area independently. We found no significant interaction effect for the two factors in CA1 and (F(1,117)= 0.08, p=0.78) and significant effect on learning condition (F(1,117)=6.59; p<0.05) and trial outcome independently (F(1,117)=5.05; p<0.05). In PFC both trial outcome (F(1,92)= 12.05, p<0.001) and learning (F(1,92)= 18.13, p<0.001) showed significant main effects, but there was also a significant interaction between factors (F(1,92)= 4.93, p<0.05). Shaded areas and errorbars denote SEM. */#: p<0.05; ***/###: p<0.001.

We found that, during learning sessions, the predicted time of the US temporal sequence strongly concentrated at the beginning of the stimulus presentation, where CS+ temporal sequences also happened (Figure 3A). This effect was strongly reduced in overtrained sessions in both CA1 and PFC, both in the general US average decoding performance (Figure 3A; rightmost panel) and in the temporal echo computed as the average performance on the main diagonal of the echo window (Figure 3B).

Because temporal echo strength decreased with learning, we hypothesized that echos could represent error signals arising during incorrect trials. Since after learning the animal committed less mistakes, that would explain weaker temporal echoes in overtrained sessions. To investigate this, we separated trials into correct (i.e., CS+ trials with anticipatory licks during trace or CS− trials with no licks) and incorrect (i.e., CS− trials with licks during trace period and CS+ trials with no licks). We then computed the time-decoding matrices (trained on all CS+ trials) for correct and incorrect CS+ test trials (Figure 3C-D). We found that incorrect CS+ trials showed a stronger US echo (in CS) compared to correct trials. This was true for learning sessions, in which the temporal echo was evident (Figure 3C,E), but also in overtrained sessions (Figure 3D,E), partially recovering the effect we found before learning. Temporal echoes were stronger on incorrect trials (two-way ANOVA with factors outcome and learning condition, main effect (outcome) CA1: p<0.05; PFC: p<0.001) and during learning sessions (main effect (learning condition) CA1: p<0.05; PFC: p<0.001). Moreover, in PFC those two factors showed a significant interaction (outcome and learning condition; p<0.05) (Figure 3E).

Interestingly, the US echo in the CS+ was stronger in incorrect trials compared to correct ones. Computing the same analysis using CS− trials (US echo in CS-) showed no difference between correct and incorrect trials S3. Thus, CS and US representations of time are initially similar, but progressively decorrelate with learning. Because CS+ and US sequences recovered an increased similarity during incorrect trials (i.e., when the animal didn’t expect the reward), we further hypothesized that the temporal sequences found were a general mechanism evoked by unexpected events. We reasoned that, although CS+ and US could initially be both perceived as unexpected events, after learning US occurrences would be predictable by CS+, leading to a change of the US-evoked temporal sequence and a decreases the similarity between the two sequences. To test this hypothesis we investigate the similarity between CS− temporal sequences and CS+ and US before and after learning.

To do so, we first used a Bayesian decoder trained to predict CS− time (as in Figure S2) to decode samples of CS+ activity. We reasoned that, if the same neural representation of time was shared between the CS+ and CS− conditions, then a classifier trained to discriminate time in CS+ trials would generalize well when tested on CS− trials; and vice versa. Alternatively, a failure of the classifiers to generalize between conditions would suggest the presence of different time encoding schema. This cross-stimulus decoder revealed that CS− and CS+ temporal sequences have a shared temporal representation, with the CS+ temporal sequence being correctly decoded during CS− in both CA1 and PFC (Figure 4A, see also Figure S4 for CS− sequences during CS+ trials). In CA1, this shared representation increased with learning, probably reflecting the strongest temporal CS+ representation in that area after learning. Moreover, the CS− temporal sequence also echoed on US temporal sequence in the test trial (Figure 4B). This effect could also be seen in the analogous case of testing CS− trials in a model trained with CS+trials, where the CS− temporal sequence was echoed in the US epoch of the CS+ decoder (Figure S4A). Similarly to CS+, the CS− temporal echo decreased after learning of the task (Figure S4B). Together those results corroborate the idea of an initial unspecific representation of time evoked by unexpected events, which acquires stimulus specificity upon learning.

**Figure 4:**
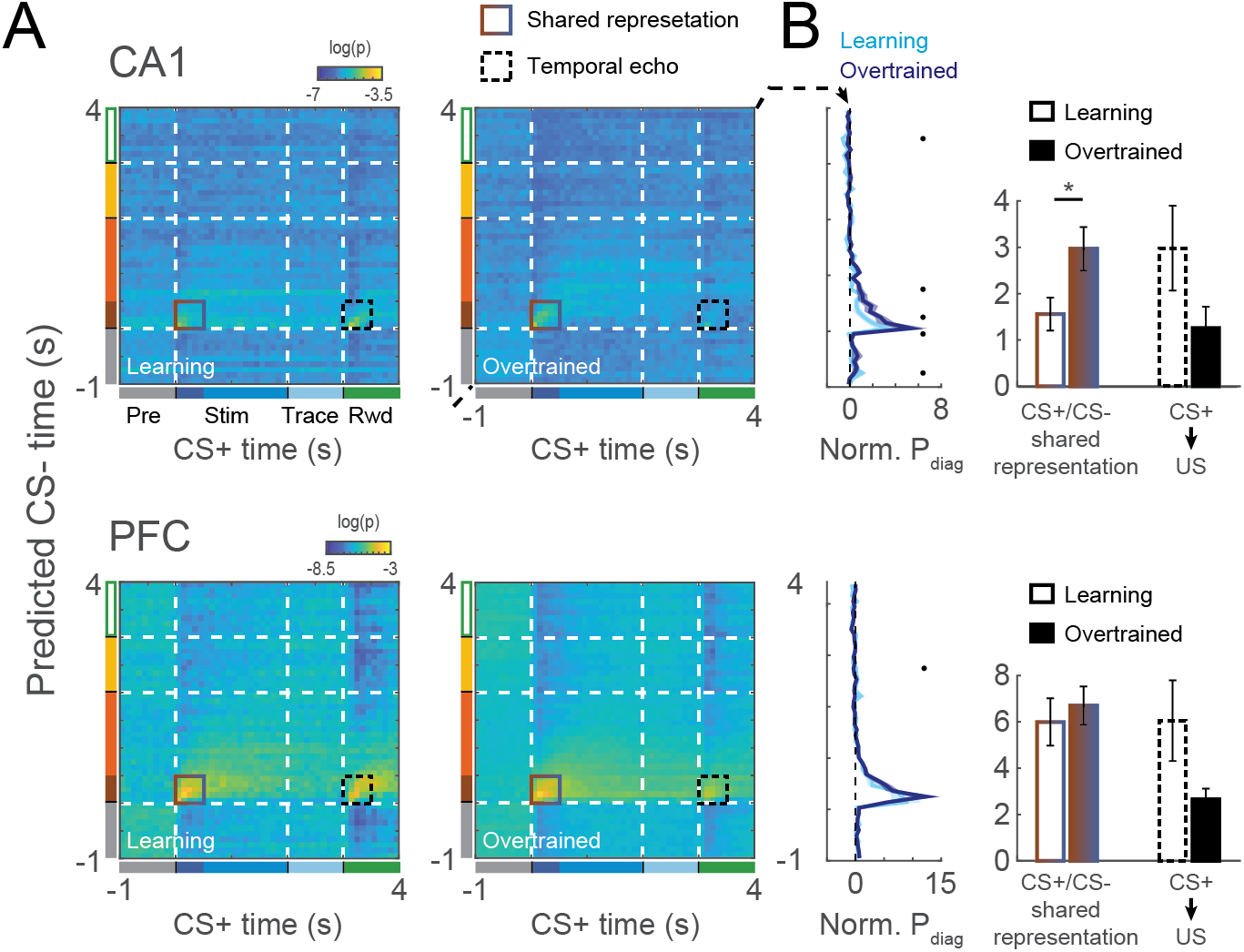
CS- have a shared temporal representation with CS+. **A.** Time-decoding matrices trained using CS− activity and used to decode time in CS+ trials in CA1 (top) and PFC (bottom). Solid boxes show areas in which a shared temporal representations can be seen. Dashed boxes show US temporal echoes on CS-. **B.** Normalized decoding performance in the main diagonal (left) and average decoding over the shared and echoed representations (right) in the matrices in A.

### Learning induces persistent stimulus representations in PFC but not CA1

We next examined whether population activity encoded stimulus identity (i.e., CS+ or CS-). We trained an ensemble of SVM classifiers to discriminate between CS+ and CS− conditions in each bin. For each classifier trained on a particular time bin, we then predicted the trial identity (CS+ or CS-) of all other bins. We used different combinations of training and testing time bins to build a temporal generalization matrix (King and Dehaene (2014); Figure 5A), a framework which allowed us to examine the temporal nature of the stimulus representations in CA1 and PFC (Figure 5A). We tested three alternative scenarios: i) A high decoding accuracy restricted to a tight region around the main diagonal, suggesting a reliable yet time-varying stimulus representation (such as stimulus-specific sequential activity); ii) An extended area of high accuracy covering a large diagonal block, suggesting a persistent and stable stimulus representation; iii) No area of significant decoding accuracy, suggesting the absence of a reliable stimulus representation (for example, in a scenario of stochastic reactivation of stimulus-specific patterns at random times in each trial). We found very different encoding dynamics in CA1 and PFC. In CA1 decoding performance seemed to concentrate near the main diagonal, and we found a significant increase in decoding performance during trace period only (Figure 5C-D). On the other hand, a strong and persistent stimulus representation encompassing both stimulus and trace periods emerged in PFC after learning (Figure 5C-D). Both effects could also be seen by looking at stimulus representation at specific time points (i.e., rows of the decoding matrices in Figure 5C), as summarized in Figure 5E. Stimulus encoding increased strongly with learning in PFC during both the stimulus and trace periods. In CA1, stimulus encoding was weaker and mildly improved with learning only during the trace period.

**Figure 5:**
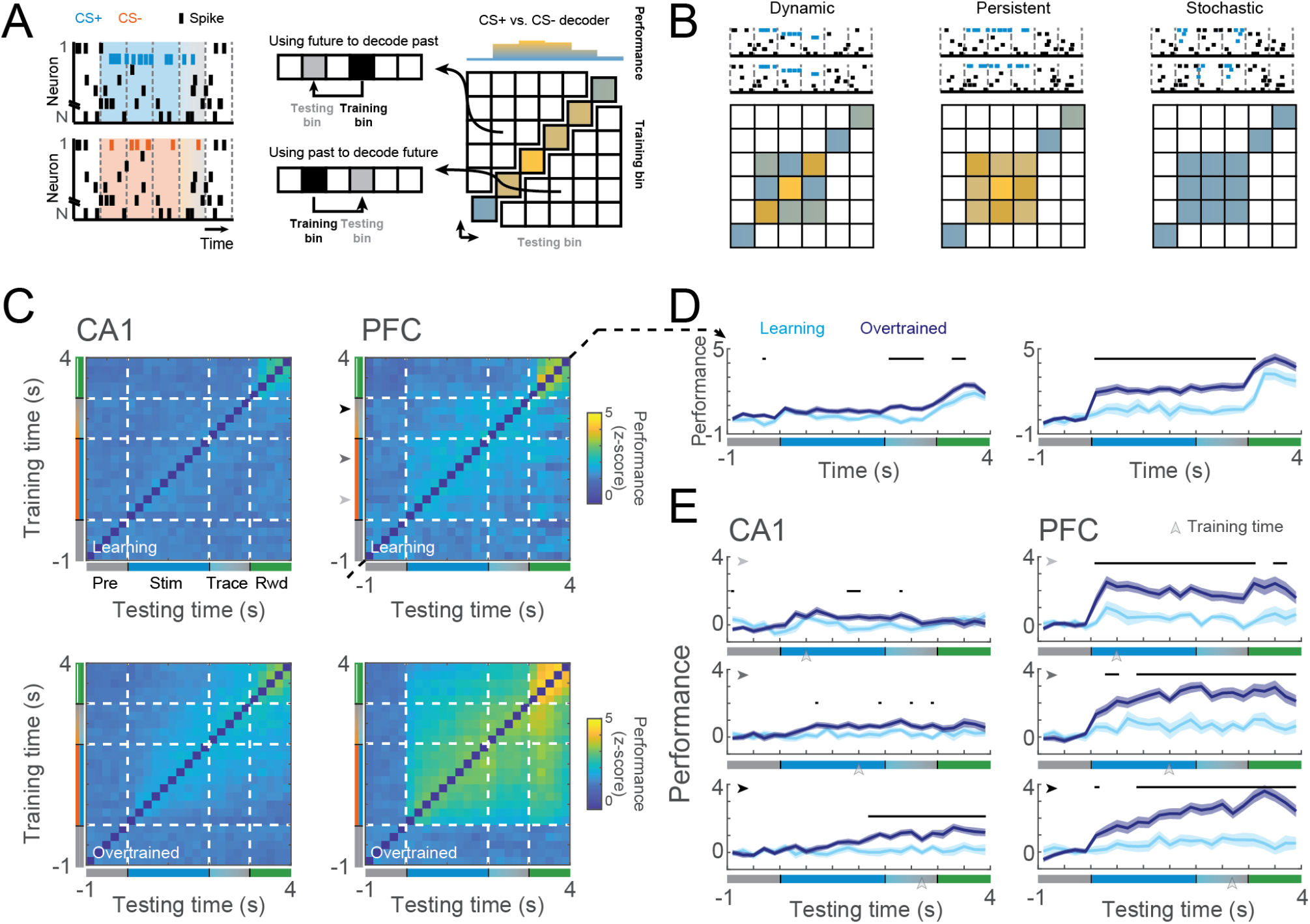
Different trial type decoding dynamics in CA1 and PFC. **A.** Schematic representation of trial type decoding matrices. The activity in each time bin was used to train a CS+/CS- SVM classifier. Test bins were evaluated on decoders based on past, current and future time bins, enabling an evaluation of stability of neural representations over time. **B.** Possible modes of neural representations. Trial decoding matrix can reveal whether the nature of the stimulus representation is time-varying (left), persistent (middle) or undetectable (right). In case of a time-varying representation, population activity would encode stimulus identity in a stereotypical trajectory, with non-generalizable decoders over the trial time. Alternatively, in case of a persistent representations, population activity would reach a fixed stable attractor, with decoders being generalized over the trial time. At last, the stimulus decoding matrix could detect no reliable stimulus representation. **C.** Mean CS decoding matrices for learning (top) and overtrained (bottom) sessions of CA1 and PFC. **D.** Average performance of the main diagonal in the matrices in C. Black dots represent bins in which learning and overtrained performance are significantly different (p<0.05; Wilcoxon ranksum test). **E.** Rows of the CS decoding matrices in C. Black arrows denote the time bin used to train the models (y-axis in C). Notice the stability of PFC stimulus representation over the stimulus and trace period while CA1 representation mildly strengthens during trace. Black traces denote different learning and overtrained performances (p<0.05; Wilcoxon ranksum test).

### Transient co-activation of neural assemblies underlies stimulus-coding neural representations

The classification analysis discussed above showed that CA1 ensembles do not encode stimulus information in a time-ordered representation consistent across trials. Alternatively, stimulus-specific assemblies could stochastically reactivate for brief intervals, at random times in different trials. To investigate the presence of transient yet stochastic reactivation of neural assemblies, we fit a hidden Markov model to each recording session, separately in CA1 and mPFC. The HMM is a probabilistic generative model for neural population data, capturing both the network organization in a small set of stereotyped neural states, each one representing epochs where neural populations fire at an approximately constant rate; and also the dynamical transitions between these states forming stochastic sequences. Figure 6A shows an example of neural activity in a CS+ trial in PFC, where multiple states were detected. In the following analyses, unless stated otherwise, we used the most-likely state to define state activation.

**Figure 6:**
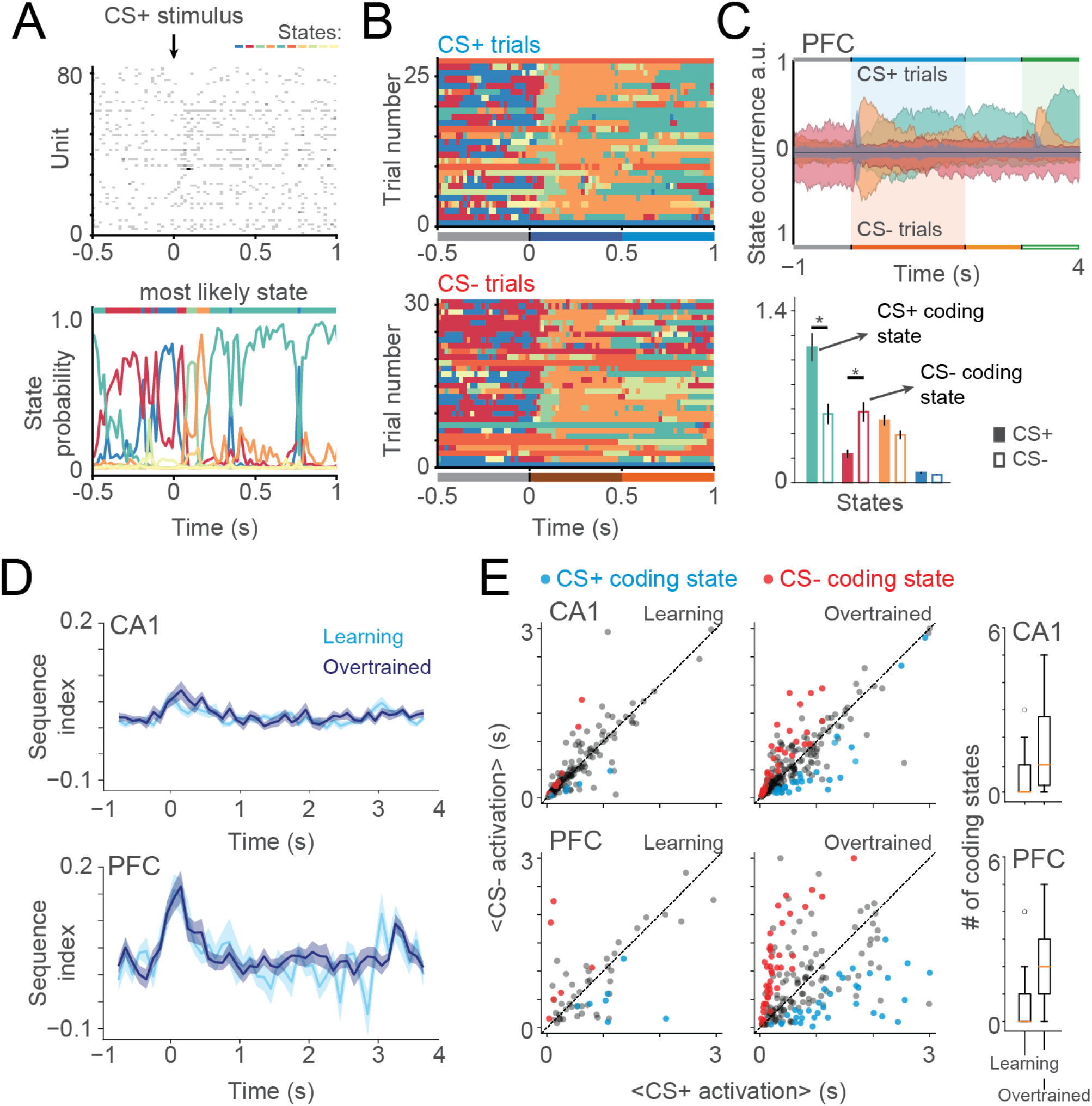
Transient co-activation of neural assemblies captured by hidden Markov models. **A.** Neural activity during a CS+ trial (top) and its respective hidden states probabilities (bottom) during an overtrained example session in PFC. Color-coded curves represent posterior probabilities of each state. At each time point, the most likely state (colored bar on top) was defined as the active state. HMM states were computed separately for CA1 and PFC. **B.** Example plot of state activation in PFC over the beginning of different CS+ (top) and CS− (bottom) trials for the session in A. Notice the visible sequential activation of states after stimulus onset. **C.** (Top) Probability of state activation over CS+ (upper quadrant) and CS− (lower quadrant) trials for some states of the session in B. (Bottom) Bar plots show the average state activation probability during stimulus and trace period. Note that some states are significantly more active CS+ (CS+ coding states) or CS− (CS− coding states). Errorbars denote SEM (*p<0.05; Wilcoxon signed rank test). **D.** State sequence index computed in a 0.5 s sliding window for learning and overtrained sessions in both areas. Notice that sequence index peaked after stimulus onset, showing that HMM states could also capture the temporal sequences dynamics. For this analysis both CS+ and CS− trials were used. **E.** Average CS+ and CS− activation during trace and stimulus period for every state in learning (left) and overtrained (middle) sessions, together with boxplots showing the distribution of coding states per session (right). Color denotes CS+ (blue) and CS− (red) coding states. Note the stronger increase in the number of coding states for PFC.

A pseudo-colored rastergram showing state activation in CA1 and PFC for trials in a representative session can be seen in Figure 6B. In general, CA1 showed a higher number of states than PFC in both learning and overtrained sessions (Figure S5B; model selection for the number of states was obtained via the Bayesian information criterion (BIC), Supp. Figure S5A; Recanatesi et al. (2022)) and state duration in CA1 was also shorter than in PFC (Figure S5C). Moreover, the probability of most-likely state in CA1 was generally lower, peaking around 0.65, than in PFC, which had usually values above 0.8 (Figure S5D,F). This suggests that PFC state activation is univocal, with no competition across states, while in CA1 states may be coactivated or partially overlapping with other states. Following our previous results, we hypothesised that HMM states would also be able to capture the presence of the temporal sequences in the form of sequential state activation. Notice that a sequential state activation can actually be seen in the example sessions in Figure 6C, in the blue, orange and green states. In order to measure that, we used a sliding window to compute the (state) sequence index - an unbiased mutual information between the state activation and their ranks across multiple trials (see Materials and Methods and Figure S6). We found a strong increase in the sequence index near the stimulus onset for PFC and a smaller peak in CA1, consistent with the temporal sequences found using the time-decoding matrices (Figure 6D). Moreover, the sequential structure was consistent across learning and overtrained periods, suggesting that the encoding of temporal information is independent of learning.

### Transient co-activation of neural assemblies underlies stimulus-coding neural representations

To investigate whether HMM states encoded stimulus identity, we used the distribution of state activation during stimulus and trace periods in CS+ or CS− trials (Figure 6C). States that were significantly more active during CS+ or CS− were defined as ’CS+ coding states’ or ’CS- coding states’, respectively (Mazzucato et al. (2019)). Before our learning criteria, only a small number of coding states were detected both PFC and CA1 (Figure 6E). Both areas showed a significant increase in the number of coding states after learning, with a larger fraction of coding states in PFC compared to CA1, despite the larger number of HMM states detected in CA1 overall and consistent with the larger stimulus-decoding accuracy found in PFC compared to CA1.

We then focused on coding states in overtrained sessions, and looked at which moments those states were most active along the trial (Figure 7A). We found that CS+ coding states increased their activation during CS+ trials, while keeping similar levels of activation in CS− trials in comparison to pre-stimulus period. Notably, the average CA1 activation of CS+ coding states in CS+ trials peaked in the middle of stimulus period, while in PFC those coding states had a sharp and steady increase in their activation levels. For CS− coding states the greatest change was also present during CS+ trials, in which the coding states were strongly inhibited, following a similar temporal dynamic in CA1 and PFC as in the CS+ coding states. Altogether, those results suggest that both CS+ and CS− coding states are originated by changes in state activation during CS+ trials. In order words, CS− coding states are indefinitely active, and inhibited during CS+ stimulus instead of being more active during CS-. This suggests that encoding stimulus identity by coding states in PFC is biased towards the relevant stimulus of the task (i.e., CS+). These results explain the single-trial origin of the poor decoding in CA1 and the strong and stable decoding in PFC discussed above. The fact that we can find coding states in CA1 and that they do not follow a stable temporal pattern of activation corroborate the hypothesis of stochastic coding states (i.e., not time-locked with the stimulus) dominating CA1 activity.

**Figure 7:**
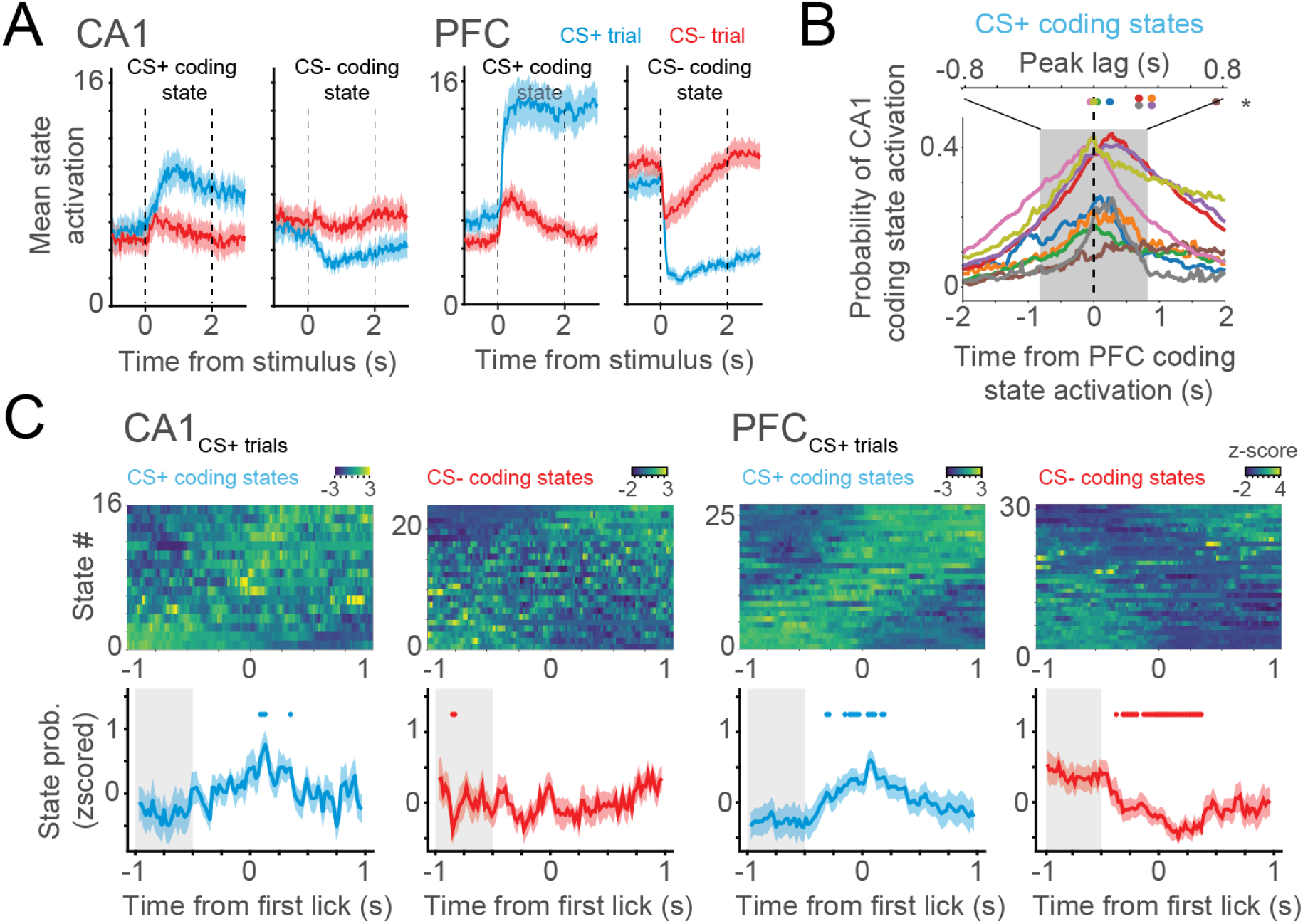
Emergence of CS coding states after learning. **A.** Average of CS+ and CS− coding states activation in both areas. The activation of coding states are shown during CS+ and CS− trials. Note that in both areas CS− coding states show increased activation during pre-stimulus period, being inactive during stimulus in CS+ trials. On the other hand CS+ states have low activation during pre-stimulus period, peaking after the stimulus presentation. **B.** Cross-correlation between CA1 and PFC (CS+) coding states activation in sessions with simultaneous recordings. Most of CA1 coding states peaked their activity after PFC coding states activation. **C.** Coding state’s activation during overtrained sessions, centered on the first lick of (CS+) trials. State activation is shown for both CS+ and CS− coding states of CA1 (left) and PFC (right). Notice that activation of CS+ coding states precedes the first lick of the trial in PFC. Similarly, CS− coding states are inhibited before the lick onset. This can not be explained by proximity with stimulus onset (see methods). Dots represent probability of state activation significantly different than the average probability in the shaded rectangle (p<0.05; Wilcoxon sign-rank test. Shaded areas denote SEM.

We next hypothesized that coding states in both areas could have an increased co-activation pattern. We then computed the cross-correlation between pairs of CS coding states simultaneously recorded in CA1 and PFC during stimulus and trace period (Figure 7B). We found that CS+ coding states in CA1 and PFC tended to co-activate, with PFC states preceding CA1 ones. Interestingly, when looking at the co-activation of CS− coding states in the same period we could see no area leading the state activation (Supp. Figure S8B). Despite those results suggesting some interplay between CS+ coding states in both areas, we found that the peak correlation between coding states was not higher than between non-coding states (Supp. Figure S8C). Moreover, the mutual information between state activation in CA1 and PFC actually decreased during the late stimulus and trace periods of CS+ trials (Supp. Figure S5G). Together, those results suggest that CS+ coding states in PFC might have a leading role into representing the trial type, but there is no task-related increase in CA1-PFC communication.

### Onset of stimulus-coding neural states predicts behavioral performance

We next investigated whether one could reliably predict the animal’s behavior in single trials from the specific onset of PFC stimulus-coding states. We hypothesized that the activation of a CS+ neural representation could lead to the behavioral expression of the conditioned response. Specifically, we examined whether the onset of a CS+ coding state in a correct CS+ trial preceded the onset of the first lick and likewise, whether the offset of CS− coding states preceded lick onset (Figure 7E). In order to do that, we computed the (z-scored) coding-state probability centered at the time of the first lick of the trial. Significance was accessed by comparing state probability to the average baseline probability (computed in a 0.5 s window from −1 s before the lick onset). To rule out effects of stimulus onset, only first licks happening 1 s after stimulus were considered. We found that specifically in PFC an increase in probability of CS+ coding states consistently preceded lick onset in CS+ trials. Analogously, CS− coding state probability consistently decreased before lick onset. In CA1, CS+ coding states sharply peaked their probability after the lick onset, which is in accordance to the posterior activation of CA1 CS+ coding states shown in Figure 7D. Crucially, PFC coding states showed similar dynamics in (erroneous) CS− trials (Supp. Figure S8A), with CS+ coding states with a tendency to peak before lick onset, and CS− coding states significantly inactivating before the first lick. Notably, this effect could not be explained by delays in lick onset detection, since the minimum inter-lick interval happened on a shorter scale (Supp. Figure S7E). We thus concluded that the transient activation of stimulus-specific neural assemblies in PFC predicts the behavioral expression of the conditioned response in single trials.

## Discussion

In this work we elucidated how the dynamics of stimulus and temporal coding in CA1 and PFC are shaped by learning during an auditory trace conditioning task. Those results are summarized in Figure 8. We found that both areas encode time via temporal sequences, evoked by stimulus and reward delivery. Interestingly, we found evidence that stimulus and reward sequences may represent surprise signals, whose representations are similar in naive animals, and progressively acquire specificity during the course of learning. We found that learning induces the emergence of a persistent and stable stimulus representation after learning in mPFC, bridging the gap from stimulus and to reward across the trace period. Looking for stimulus representation on neural states in single-trials confirmed most of our previous results and further showed that transient activation of stimulus-coding states in mPFC reliably predicted the onset of the first lick response, providing a link between neural coding and behavior.

**Figure 8:**
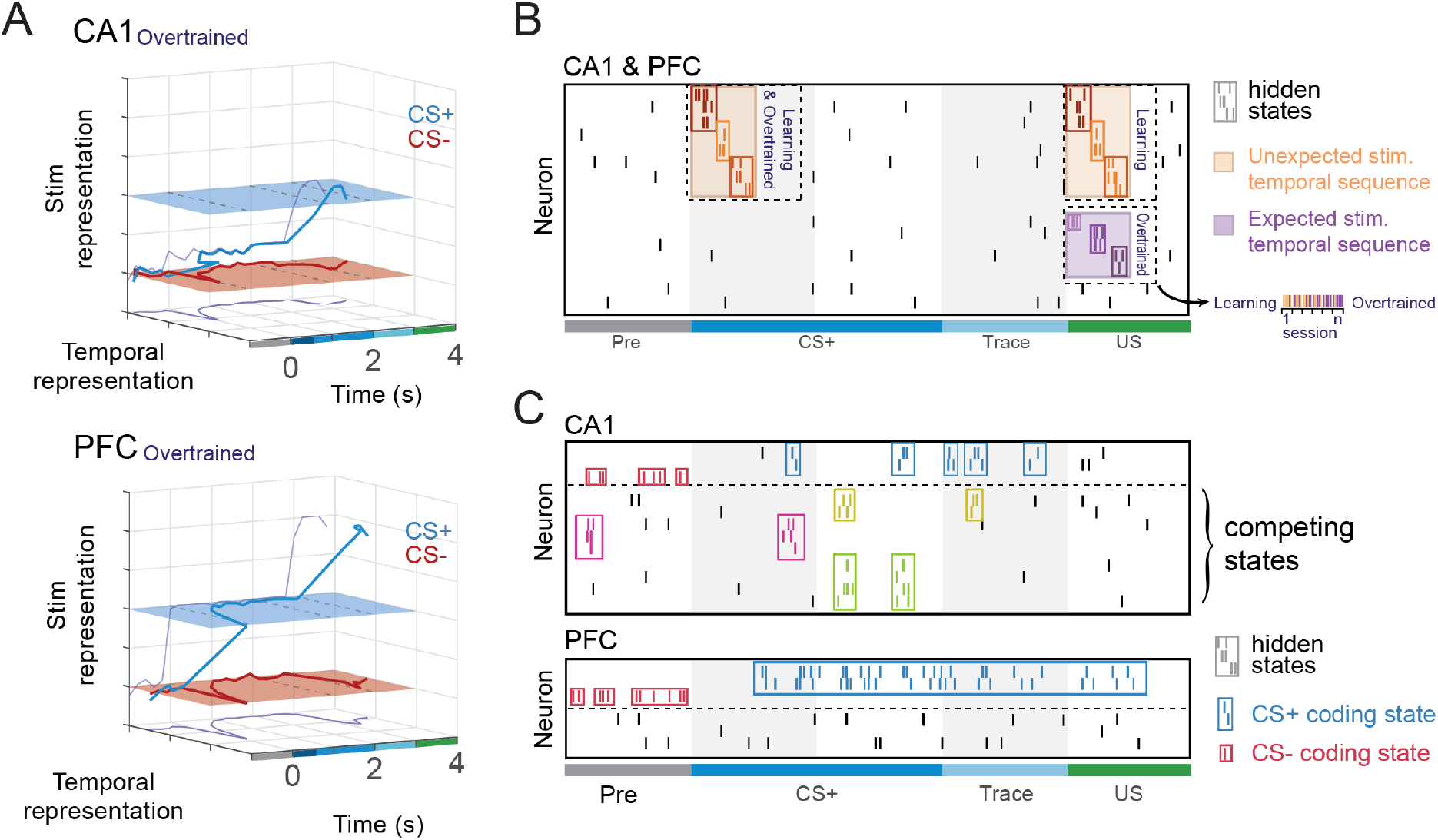
Schematic representation of temporal and stimulus coding in CA1 and PFC A. Joint temporal dynamics of time and stimulus decoding in overtrained sessions. In CA1 overtrained sessions the stimulus representation of CS+ and CS− lie most of the time in the same plane, starting to increase during trace, and peaking at reward consumption, where CS+ and CS− are differentiable. In contrast, PFC stimulus representation of CS+ differentiates from baseline and CS− representation right after stimulus onset. This effect was also accompanied by a peak on temporal representation, present after CS+ and CS− onset on both areas. The temporal representation also peaked after US onset. **B.** Learning dynamics of temporal sequences and hidden states in both CA1 and PFC. Activation of hidden states captured by HMMs are shown in the small colored boxes. Unexpected stimulus triggers the onset of a generic temporal sequence that can be captured by the states (orange boxes). After learning, the triggered sequence of an expected event (US) changes, disentangling from the unexpected sequence (purple box). Note that, even after learning, error trials (when the US is again unexpected) can recover the activation of unexpected temporal sequences (horizontal colorbar). **C.** Coding state dynamics in CA1 (top) and PFC (bottom). CS+ coding states (blue) in both areas are mostly active during CS+ stimulus presentation, while CS− coding states (red) are indiscriminately active, but inhibited during CS+. In CA1, coding states compete with other states in a way that probability of the most likely state is generally low. On the other hand, there was almost no competition among PFC states. Moreover, decoding performance was strong and generalizable in PFC, suggesting persistent activation of CS+ coding states. In contrast, the milder, time-specific decoding performance in CA1 can be explained by a transient and stochastic activation of coding states, that competed with other states.

### Time encoding and surprise signals

Since time is an important component of declarative memory, and given the role of the hippocampus in this type of memory, it’s not surprising to find some form of temporal coding in this area. In fact, hippocampal neurons can have timely coordinated spikes in the form of phase precessing place cells and time cells (Eichenbaum (2014), O’Keefe and Recce (1993)). The increase in temporal decoding we found in CA1 upon learning indicates that the learning process sculpts the transient response to the CS in a behavior-dependent way, making it more temporally unique, and longer-lasting (Figure 2). It has been shown that the hippocampus (Solomon et al. (1986), Bangasser et al. (2006)), and specifically CA1 (Kesner et al. (2005)), is needed to create the CS-US associations in trace conditioning tasks. In accordance with this, previous works suggest that time cells may bridge stimulus representation during the trace period (Pastalkova et al. (2008b), MacDonald et al. (2011, 2013), Kitamura et al. (2015), Sellami et al. (2017)). Although we found temporal representations of the stimulus, differently from those other works, they faded after the first 0.5 s of stimulus presentation (Figure 8A). This is similar to other CA1 studies showing no time encoding in calcium traces bridging CS and US during trace fear conditioning (Ahmed et al. (2020)). In our case, our shorter time-scale analyses allowed us to extend those findings and find the fast onset and offset of temporal sequences. This might indicate a lower hippocampal load in this particular task and an CA1-independent maintenance of stimulus representation during the trace period, as manifested in mPFC.

Interestingly, the increase in time encoding with learning was seen specifically after CS+ presentation - the relevant stimulus of the task - and not after CS− or US (which also presented high values of temporal decoding, but no changes after learning) (Figure 2C; see also Figure S2). It has been previously shown that CA1 is important to activate cortical memory representations during memory retrieval (Tanaka et al. (2014)). It might be that the emergence of a stronger temporal sequences is the mechanism used to process the sensory information arriving in CA1 from the entorhinal cortex and propagate them to the PFC, where the stimulus-context associations are held (Euston et al. (2012)).

Moreover, before learning CS temporal sequences seemed to be echoed during the US presentation indicating that initially the same temporal pattern was elicited after all the three events (CS+, CS− and US) (Figure 3). After learning, CS and US similarity drastically decreased in both areas, except during putative incorrect trials where their pre-learning similarity was partially recovered (Figure 3). This suggests that, in error trials where the CS+ stimulus identity was likely not encoded and thus the animal did not expect a reward, US-evoked sequences may encode a surprise signal, more similar to the CS-evoked sequence, as it occurred before learning. Additionally, this effect seems to be supported by changes in US temporal sequence, while CS+ and CS− shared representation increases with learning (Figure 4).

Together those results suggest a common temporal component triggered by surprise. For example, it might be that initially unexpected events (CS or US) evoked a partially overlapping temporal sequence, whose shared element represents a ’surprise’ signal. After learning, the surprise element wanes from the US since it then becomes expected (putative correct trials), leading to the emergence of stimulus-specific responses and enabling a decorrelation between CS+ and CS− (Figure 8B) and between those and the US. It is worth to note that, despite the difference between correct and incorrect trials in US echo in CS+, there was no such difference for the US echo in CS− (Figure S3). This suggests that US temporal sequence are differently captured by CS+ and CS− decoders, and there might be an additional component beyond ’surprise’ playing a role in the similarity between CS+ and US.

### Stimulus coding via mPFC persistent activity

Even tough temporal sequences encoding the outcome signal of the task (i.e., the presence or not of the US) are present in both CA1 and PFC at the end of the trial, those two areas show different stimulus encoding profiles. While in overtrained animals CA1 decoders only had a mild increase in performance during trace period, mPFC showed a strong and stable stimulus decoding since CS presentation (Figure 5 and 8A). Thus, beyond the outcome of the task, mPFC neurons also contained information about the stimulus, even in its absence, which is in accordance to previous works (Asaad et al. (2017)) and further support the idea that mPFC holds the stimulus-action maps.

mPFC stimulus representation could be generalized from stimulus to trace period, suggesting the neuronal dynamic encoding the stimulus in this area reach a stable attractor state. On the other hand, CA1 decoders could not be well-generalized for different time periods. Could the poor decoding performance in CA1 be explained by fast stimulus-specific patterns reactivated stochastically at random times (i.e., not time-locked with the stimulus)? A single-trial pattern analysis based on hidden Markov models managed to reveal stimulus-coding states in both CA1 and PFC, giving support to the hypothesis of stochastic reactivations in CA1 (7). Moreover, it might be that in mPFC assemblies compete with each other in a winner-takes-all dynamic, with the strongest assembly dominating the state of the network, while in CA1 there are different independent patterns happening concomitantly, making it hard to identify independent states (Figure 8C). This idea is supported by the lower and higher HMM posterior probabilities found in CA1 and PFC, respectively, (Figure S7D and F), and corroborates previous works showing neuronal representations encoding different context in mPFC are well separated while highly overlapping in CA1 (Hyman et al. (2012), Leutgeb et al. (2004)), and previous findings showing very sparse stimulus selectivity in CA1 (Ahmed et al. (2020)).

CA1 and PFC showed the presence of both CS+ and CS− coding states after learning (Figure 7B). Interestingly, we found that CS− coding states were unspecific to the stimulus period, being active during pre-stimulus period as well (Figure 7C). On the other hand, the occurrence of CS+ coding states was restricted to the CS+ stimulus epoch. In either case (for both CS+ and CS− coding states) coding properties arise due to a higher state modulation (activation or inhibition) in CS+ trials, the relevant stimulus for US prediction. Moreover, for PFC the activation of a CS+ coding state reliably preceded the upcoming first lick of the trial, thus ruling out that it is driven by backdrop of movements reflected in lick modulated neurons as previously reported in mPFC (Figure 7 C; see also the inter-lick interval in Figure S7E). Crucially, we found that the onset/offset of PFC coding states in single trials reliably predicted upcoming erroneous licks in CS− error trials (Supp. Figure S8). This suggests the presence of a causal relationship between transient activation of stimulus representations in mPFC and an animal decision to lick.

### CA1-PFC coordination

We did not find evidence for general coordination between PFC and CA1 states. The peak cross-correlation values of CS+ coding states were not significantly different than for CS− coding states and even non-coding states (Supp. Figure S8C), reinforcing the idea that there is no task-dependent increase in CA1-PFC coordination. In addition to that, there was no increase in the mutual information between CA1 and PFC states over the task (Supp. Figure S5G). It seems that the presence of coordinated activity between those two areas may vary according to the type and period of the task (e.g., sleep, sharp-wave ripples, sensory encoding), which would explain previous work showing both dependent Jadhav et al. (2016), Peyrache et al. (2009), Shin et al. (2019) and independent (Kaefer et al. (2020), Klee et al. (2021)) co-activation of neuronal patterns in those areas. In our case, any relevant interplay between those two areas during the late stimulus or trace period could not be captured by HMM states. However, this is also consistent with a scenario where CA1 encodes representations of multiple task-variables in parallel or in a temporally continuous rather than discrete way. An HMM analysis would likely fail in these scenarios, and alternative methods such as factor analysis (Yu et al. (2008)) or matrix factorization (Mackevicius et al. (2019)) might help overcome these issues.

Despite the lack of coordination between states in the two areas, the cross-correlation between coding states in both ares showed that PFC leads CA1 in the activation of CS+ coding states, showing the importance of our single trial analysis with the HMMs. In fact, this late activation of CA1 (CS+) coding states could also be seen when investigating states activity around the first lick of the trial. For CA1, CS+ coding states activation peaked later, after the first lick of the trial. This raises the possibility that, while CS+ coding states in PFC are involved with the actual decision to lick, in CA1 they reflect the sensory signal arising from the entorhinal cortex after the licking behaviour.

In summary, our findings unveiled rich and complex dynamics in CA1 and mPFC during an auditory trace conditioning, opening new interesting questions. Our results suggests that in our trace conditioning task CA1-mPFC interplay might happen mostly following the CS presentation, when time encoding is higher in both areas (Figure 8A,B). After learning, strong, sustained and stimulus-specific representations emerged in mPFC, bridging across the trace period, with neural states predicting the animal behavior in single trials (Figure 8C).Because those temporal representations also encoded the outcome of the task, the emergence of stronger and more specific temporal sequences in CA1 might support stimulus encoding and help to organize the following stable memory representation in PFC.

## Materials and Methods

### Animals

In this study we used a total of 17 male C57Bl/6 mice. From this total, seven animals were used for (acute) silicon probe recordings in the dorsal hippocampus, six for silicon probe recordings in the medial prefrontal cortex, and four additional animals were used for simultaneous silicon probe recordings. Animals were kept in a reversed light/dark cycle, and had access to food ad libitum until two days after the surgery, when food restriction was introduced (90% of initial body weight). A detailed description of experimental procedures can be found in Klee et al. (2021). This study was approved by the Dutch Central Commissie Dierproeven (CCD) and conducted in accordance with the Experiments on Animals Act (Dutch law) and the European Directive 2010/63/EU on animal research.

### Surgical Procedures, behavioral training and data collection

Animals were kept under isoflurane (1-2%) anesthesia and placed in the stereotaxic frame. Carprofen (5 mg/kg) and lidocaine were subcutaneously injected in the area of the scalp before skull exposure. A custom-made circular head-fixation plate was placed on the skull and fixated with dental cement (Super-Bond C&B). A skull screw was placed over the cerebellum to serve as ground and reference for the later electrophysiological recordings. For animals used in CA1 recordings, a craniotomy was performed at the left hippocampus (−2.3 mm posterior and +1.5 lateral to Bregma). For animals used in PFC recordings the craniotomy was done over the left frontal cortex (+1.78 mm anterior and +0.4 lateral to bregma). A silicon elastomer (Body Double Fast, Smooth-on) was then used to covered the exposed skull until the first recording. After at least 2 days of recovery from surgery, animals began habituation to the head-fixed setup which consisted of two rods that could be screwed to either side of the implanted head-plate. In the head-fixed setup animals could move through a virtual linear track using a air-supported spherical treadmill. Feedback from treadmill movement was projected in a spherical screen surround the animal head. A plastic reward port delivering soy-milk was placed 0.5 cm aways from the posterior lip of the animal. Animals underwent at least 6 habituation sessions (3 sessions of 10 min per day) in which they received about 0.2 ml of soy milk before the beginning of trace electrophysiological recordings. The appetitive auditory trace conditioning task required the animals to associate a specific sound stimulus (the positive conditioned stimulus; CS+) to a later reward (unconditioned stimulus; US), in contrast to a second stimulus not associated with the reward (CS-). In our task, CS+ and CS− stimuli lasted for 2 s, and in the case of CS+ trials, the stimulus was followed by a silence period of 1 s (trace period) and then US delivery. Trials were then structured as 1 s of pre-stimulus, 2 s of stimulus, 1 s of trace, and 1 s of reward period. Learning of the task was accessed via anticipatory licking, detected using a infrared sensor beam in front of the reward spout. Sessions in which the amount of anticipatory licks during the trace period in CS+ trials was significantly different than in CS− trials (t-test) were classified as overtrained, while the others were classified as learning sessions. CS+ and CS− trials were presented in a pseudo-random order, and counterbalanced during the session.

### Neural data recording and analysis

At the beginning of each recording session the mice were head-fixed and had the elastomer removed for skull exposure. One or two 128-channel silicon probes were then acutely inserted above the previously prepared craniotomies using a micromanipulator. For mPFC recordings, electrodes were slowly lowered up until −2.0 mm ventral to Bregma. In CA1 recordings the electrophysiological signal was monitored for high power over the ripple frequency band (150-300 Hz), and spiking activity (600-2000 Hz). Electrodes were placed so that the highest ripple power and spiking activity were localized in the middle of the probe. Spike sorting was automatically done using Kilosort and visually inspected using ‘phy’ gui (https://github.com/cortex-lab/phy). Following analysis, unless specified otherwise, were done using custom-made Matlab scripts.

#### Decoding of trial time

Temporal coding at the single session level was assessed through the ability of decoding the time of the trial using a Bayesian classifier. For each session, we first binned trials into 100 ms bins with no overlap. The number of trials for each stimulus was balanced between conditions. A naive-Bayesian classifier was trained with a leave-one-out cross validation scheme in for each iteration a pair of CS+/CS- trial was used to test. The classifier modeled neuronal firing as an inhomogeneous Poisson process whose rate was a function of trial time and neurons were assumed to be independent given trial time.

Using test samples of a given time bin, we estimated its probability of being decoded as each of the other time bins of the trial. We then stacked the probability distributions of all the time bins into a confusion matrix (’time-decoding’ matrix), pseudo-colored with the probability of predicting a tested time bin in the x-axis, as one of the different time bins in the y-axis. Decoding probability values of each pair of time bins were z-scored using the mean and the standard deviation from a surrogate distribution (100 surrogates). Surrogates were computed by shuffling the time bin identity of each trial in the training set before computing the Bayesian decoder in each fold.

Temporal sequences could be seen in the time-decoding matrix as high decoding values concentrated near the main diagonal. We selected two windows to investigate the emergence of temporal sequences, one at the stimulus onset (0-0.5 s; 100-ms-bins) and another after reward onset (3.1-3.6 s; 100-ms-bins). Temporal sequence strength was then defined as the mean probability over the main diagonal of time-decoding matrices at those windows. Reward window was delayed 100 ms from reward presentation based on the echo of the stimulus temporal sequence, which started at this moment. This was probably due to the fact that the animal needs to lick the reward to perceive it, which can introduce a sampling delay.

To assess significance of the temporal sequences, their strength was tested against the strength of a null temporal sequence of equal size, extracted from pre-stimulus period (Wilcoxon sign-rank test). Comparison between temporal sequence strength across different areas and learning condition was done using aligned-rank transform in order to normalize non-parametric data. The equivalent of a two-way Anova was then performed (see ARTool toolbox, Wobbrock et al. (2011)).

US echo in CS+ (or CS-) temporal sequences was evaluated by analyzing the probabilities of decoding time bins of reward period (after US presentation) as time bins up to the end of trace period. This was done to avoid influences on the strength of the US temporal sequence itself on the echo estimation. Echo strength was computed in a similar way as the temporal sequence strength.

#### Decoding of stimulus identity

We performed a cross-validated classification analysis to investigate the stability of stimulus representations. Neuronal activity was divided into bins of 200 ms with no overlap and used to train an ensemble of support-vector machines (SVM) binary classifiers. Each binary classifier was trained to classify CS+ and CS− trials using the activity of a given time bin. We then used test bins from previous, subsequent, or current times to see how good the prediction could be generalized. The number of CS+ and CS− samples was always balanced and paired prior to training. Because the size of sample pool was different when training and testing in the same time bin compared to training and testing with different time bins, we used different cross-validation schemes for those two situations. In the first case (training and testing in the same time bin), we used a leave-one-out cross-validation, applied to pairs of CS+ and CS− trials. In the second case (training and testing bins different), we used the entire pool of trials to train, testing in all trials from the other time bin.

For each of those two cases we also trained surrogate classifiers (100 surrogates) in which the identity of the training trials was randomized. We then used the surrogate distribution of performance to normalize (z-score) the performance values of each classifier. Similar to the time decoding matrices, classifiers with different training and testing bins were stacked together forming a trial-type confusion matrix with training time bin in the y-axis and testing (time) bin in the x-axis. Notice that in those matrices we omit the main diagonal, that has different sample statistics and cross-validation scheme. The main diagonal is shown separately.

#### Computing hidden Markov models

To investigate whether stimulus identity was encoded by fast co-activation of neuronal assemblies, we first binned spikes in 20-ms bins with no overlap. HMMs were computed using Linderman et al. (2020) and assuming that neuron’s firing rate followed a Poisson distribution. Because during training of HMMs each trial is initialized in the same state, we included an additional second before the stimulus in those analyses to account for stabilization of states (i.e., training was done using 2 s before CS up to 1 s after US). For each session, CS+ and CS− trials were balanced and split in a 5-fold cross validation scheme. The Bayesian information criterion (BIC; Recanatesi et al. (2022)) was used to select the best number of hidden states of each model (ranging from 2 to 20). Using Akaike information criterion instead held more states, but qualitatively similar results (data not shown).

After selecting the number of hidden states, we used the best model to predict the posterior probability of each state across trials. We then defined a state as ’active’ whenever that was the most-likely state, allowing us to have a mean state activation during stimulus and trace period (Mazzucato et al. (2019, 2015)). We looked for hidden states with significantly different activation in CS+ and CS− trials during the stimulus and trace periods (p<0.05, Wilcoxon ranksum test). For this analysis data was preprocessed defining valid trials for each state as trials with at least one activation. This was done to avoid permanent state transitions due to changes in recording stability occurring across the session. To account for that, for each state only trials with at least one activation were taken into consideration. CS+ and CS− coding states were then defined as states that were significantly more active during CS+ and CS− (p<0.05; Wilcoxon signed-rank tests), respectively. This same analysis was repeated using more conservative criteria for state activation (Supp. Figure S7). We used a threshold of 0.6 for CA1 and 0.8 for PFC, accordingly to the probability distribution of states in the two areas (Supp. Figure S5D,F). There was no qualitative difference between the results using the most-likely state and the conservative thresholds.

We also computed the state activation triggered by the first lick of the trial focusing on CS+ and CS− coding states. To avoid effects driven by the stimulus, we restricted this analysis to trials in which the first lick happened at least 1 s after the stimulus onset. Also, we computed the inter-lick interval distribution of the licks pooled across all animals to have an upper bound estimation of the delay between the onset and detection of the licks (Figure S7 E).

We used the simultaneous recordings to compute the correlation and mutual information between PFC and CA1 hidden states. To compute the mutual information across the trials we first computed the most likely state of each 20-ms bin (i.e., the state with higher probability). Then, the adjusted mutual information was computed in bins of 100 ms (5 20-ms bins), separately for CS+ and CS− trials.

#### State sequence index

In order to evaluate whether HMM states could also capture the temporal sequences unveiled by the time-decoding matrices, we computed a (state) sequence index. First, for each trial we computed the activation rank of the states present in a given window (Supp. Figure S6A). State rank was defined as the order of the mean activation time within the window. The ranks of each trial were then accumulated in a state-rank matrix, from which the mutual information (MI) was extracted (Supp. Figure S6B-C). A surrogate MI distribution was computed by randomizing the ranks on each trial independently. The sequence index was then defined by the MI between state activation and rank minus the average surrogate MI (i.e., an unbiased MI). The sequence index was then computed using a 5 s sliding window. Notice that differently from other methods to investigate sequence activation, the sequence index defined here does not require a template sequence, being suited for unsupervised sequence discovery over aligned events.

## Acknowledgments

We thank M.B. Maidana Capitán, R. Pedrosa and V. Lopes-dos-Santos for fruitfull discussions and feedback in earlier versions of the manuscript. Funding was provided by a German Studiensstiftung fellowship (to JK), by the Dutch NWA ’Bio-Art’ project (to FPB), and by the NWO Top-grant no. 612.001.853 (to FPB).

## Data and code availability

The data and codes used in this study are partially reported on the Donders Repository (https://doi.org/10.34973/hp7x-4241) and available upon reasonable request.

## Author contributions

Conceptualization, J.K. and F.B.; Methodology, B.S., J.K., L.M. and F.B.; Experiments, J.K.; Formal Analysis,B.S.; Resources, F.B.; Writing – Original Draft, B.S.; Writing – Reviewing & Editing, B.S., J.K., L.M., F.B.;Supervision, L.M. and F.B.;

## Supplementary Figures and Methods

### Measuring dimensionality of temporal representations

To calculate how the dimensionality of the stimulus representation varied over the trial. For that, we used principal component analysis to computed a relative dimensionality of the populational activity for each time bin. This measure was defined as ratio between the amount of principal components that accounted for 90% of the variance in a given time bin and the respective amount computed during the pre-stimulus period. We found that both during CS+ and CS− trials, CA1 neural activity showed a decrease in dimensionality after the stimulus onset (i.e., when temporal sequences appeared) and this effect was present both before and after learning. On the other hand, PFC showed a similar decrease, but restricted to overtrained sessions (Figure S2B).

**Figure S1:**
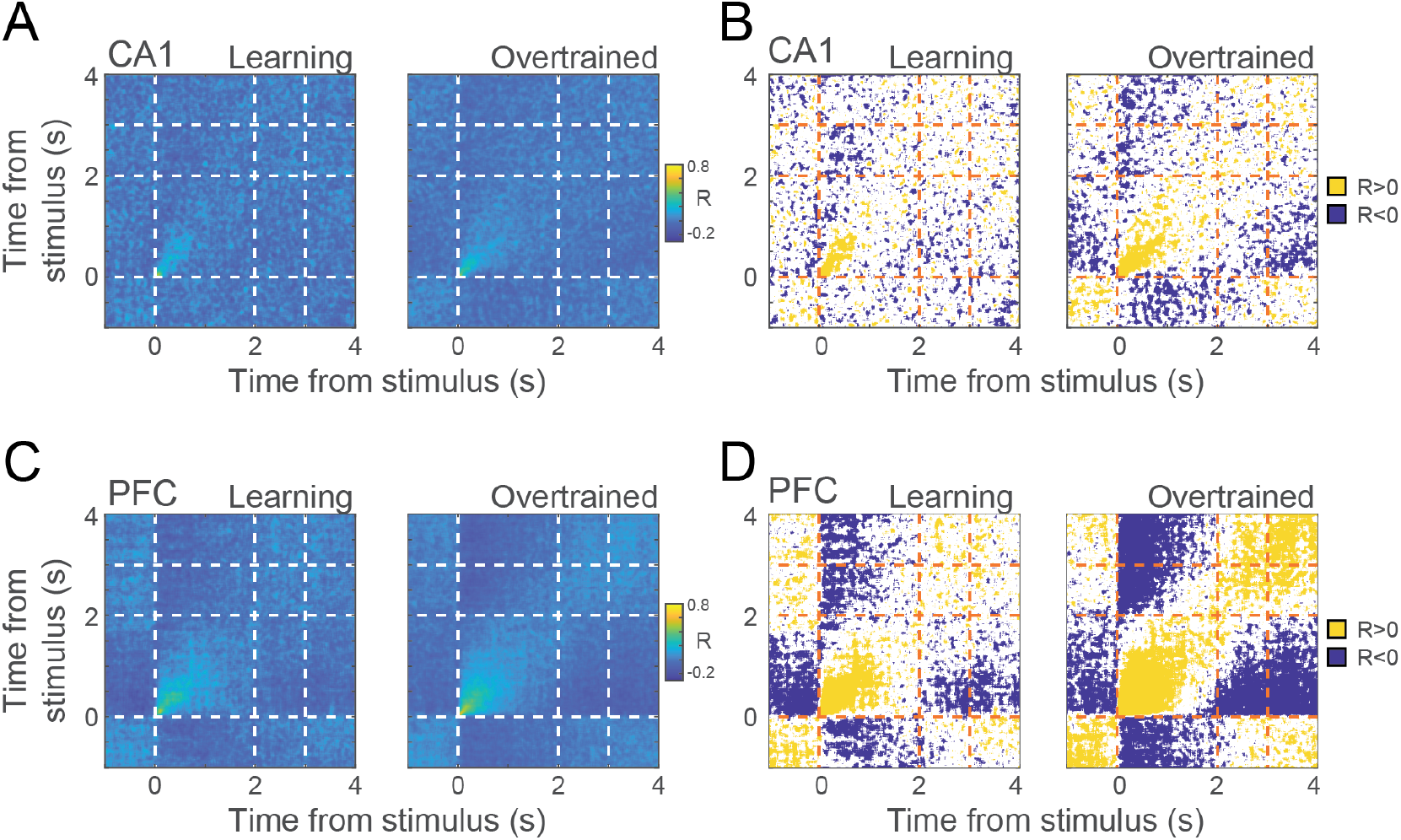
Temporal structure in pooled activity. **A.** Correlation between mean activity of all CA1 neurons on even and odd trials before and after learning (see Figure 1D). **B.** Significant correlation values (p<0.05 compared from a surrogate distribution) for A. **C** and **D** same as A and B, but for PFC.

**Figure S2:**
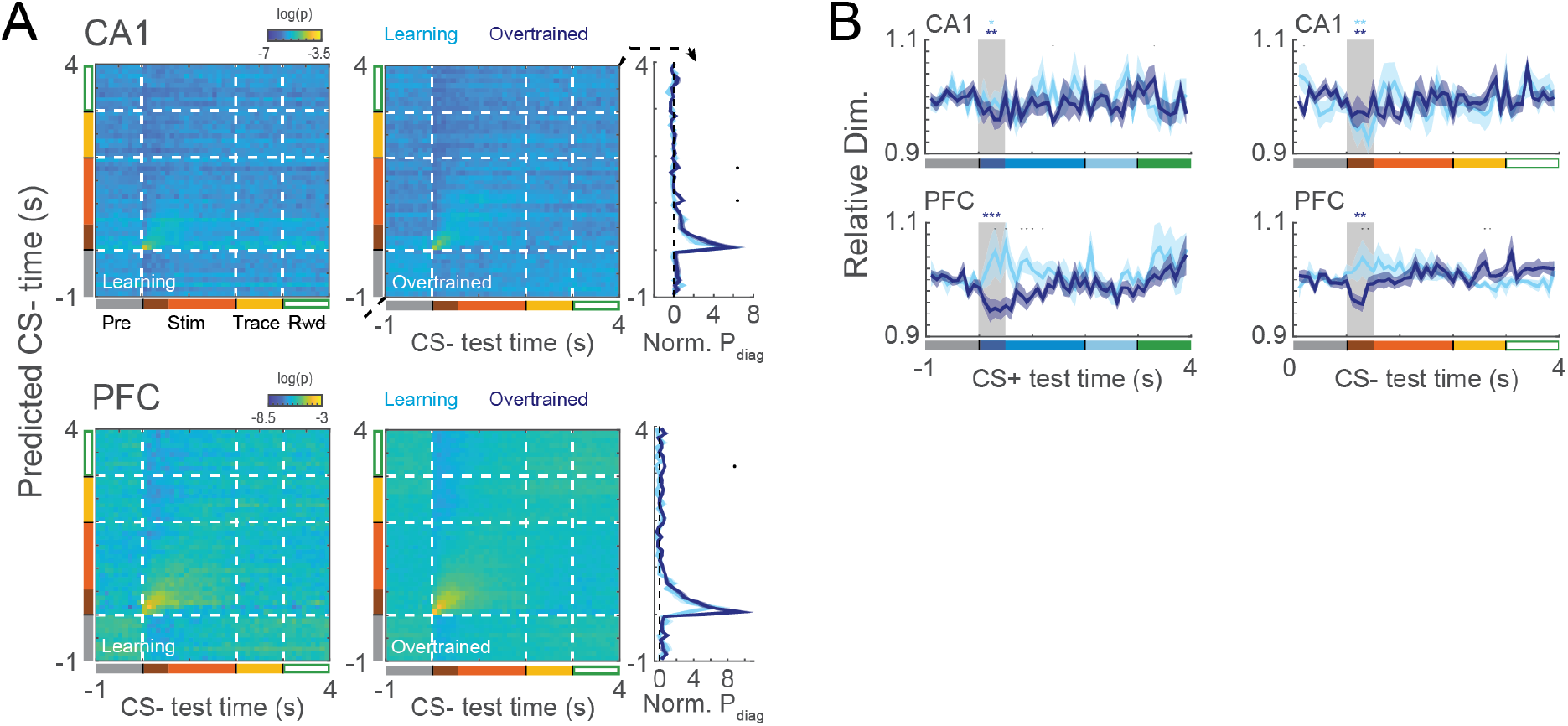
Emergence of temporal sequences in CS−. **A.** Mean time decoding matrices for learning and overtrained sessions together with the average probability on the main diagonal (rightmost panel). Decoders were trained and tested using CS− activity from CA1 (top) and PFC (bottom). Black dots denote times in which learning and overtrained decoding are different (Wilcoxon ranksum, p<0.05). **B.** Relative dimensionality of the population activity over time for CS+ (left) and CS− (right) trials. Relative dimensionality was measured as the fraction of principal components necessary to explain 90% of the data variance over each time bin. Note the lower dimensionality after stimulus presentation (in the same period of CS+ and CS− temporal representations) in CA1 before and after learning. PFC sessions showed a similar decrease only in overtrained sessions.

**Figure S3:**
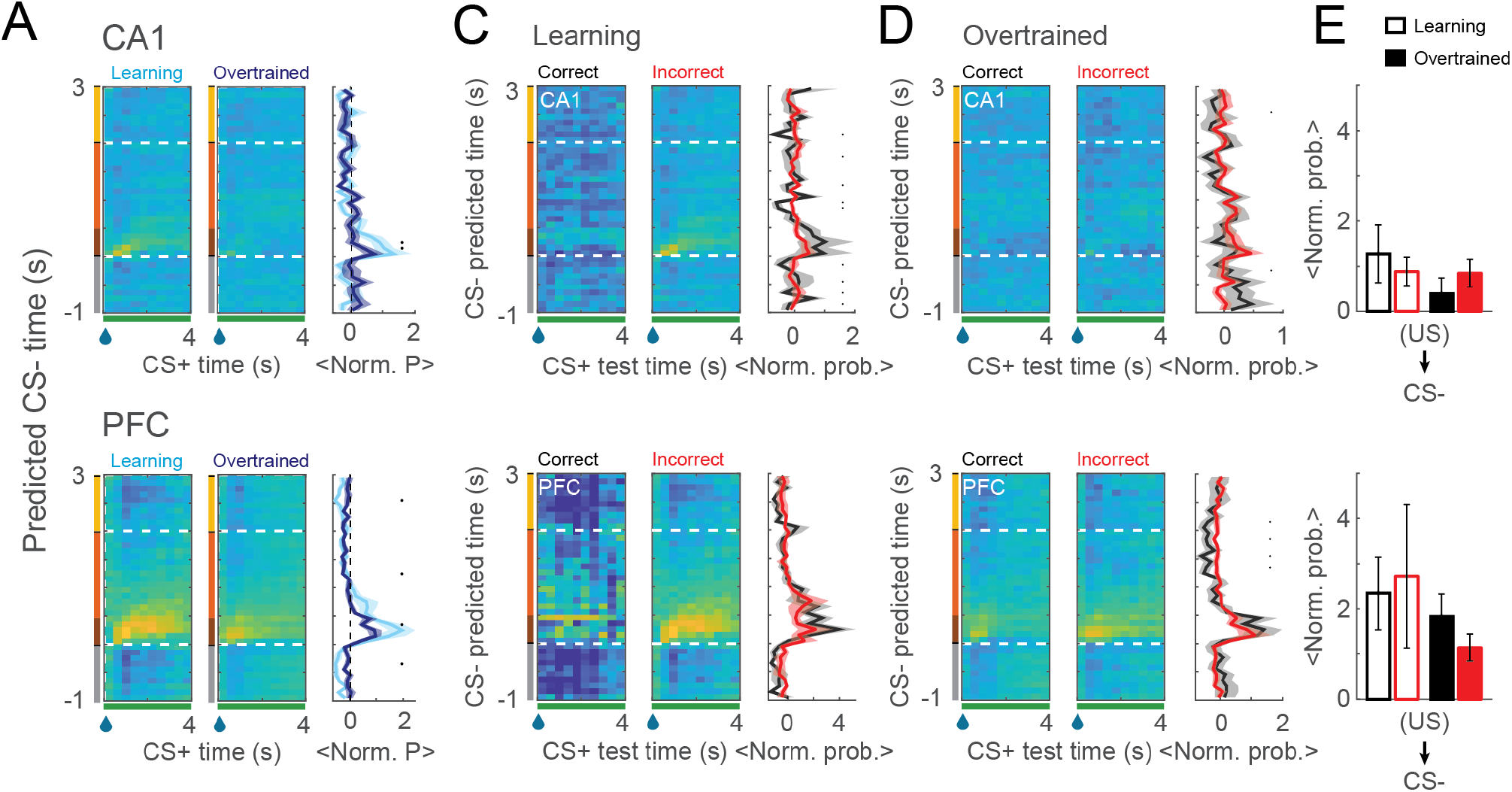
Similarity between CS− and US temporal representations is not modulated by learning or error signal of trial outcome. **A.** Similar to Figure 3, but using CS− trials for training. Time decoding matrices trained using CS− activity before the US, and used to decode US time (left and middle). Average (normalized) probabilities for the US period are shown for learning and overtrained sessions (right). **B.** Mean decoding in the temporal echo window for CA1 and PFC. **C-D.** Time-decoding matrices as in A, but showing the temporal echo in correct and incorrect trials from learning (C) and overtrained (D) sessions. **E.** Average performance on the temporal echo window (main diagonal) for CA1 (top) and PFC (bottom). Aligned rank transform followed by a two-way ANOVA using trial outcome (i.e., correct vs. incorrect) and learning condition as factors for each area independently revealed no significant interaction or main effect in either area. Shaded areas and errorbars denote SEM. Black dots denote p<0.05.

**Figure S4:**
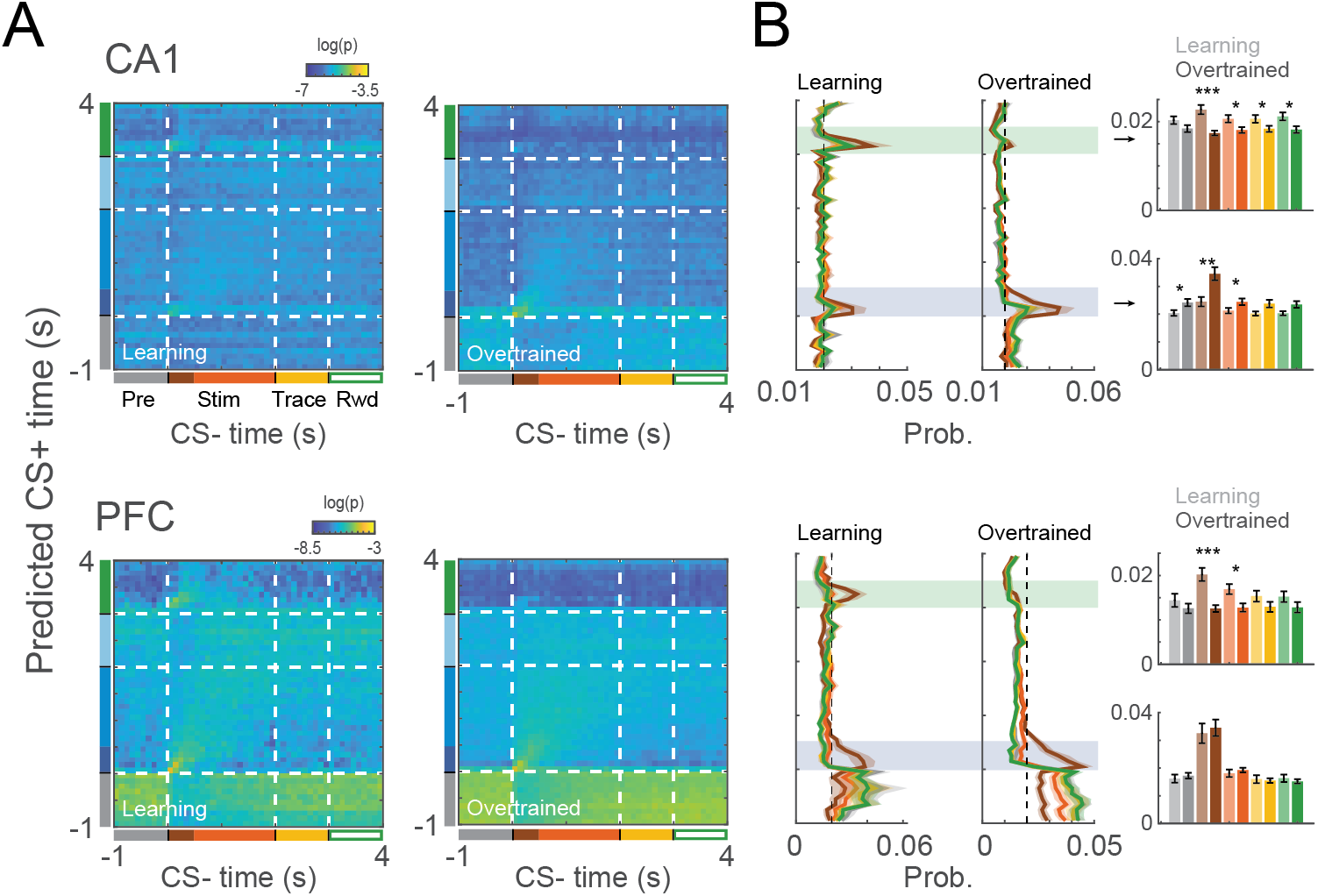
US echo in CS− before and after learning. **A.** Time decoding matrices trained using CS+ activity before the US, and used to decode US time in CS− trials during learning (left) and overtrained (right) sessions for CA1 and PFC. **B.** Average decoding probability for testing bins in different periods of the trial (right and middle), together with the mean probability in the CS and US temporal windows (shaded rectangles). Notice the decrease of similarity between CS− stimulus transient and US after learning for both areas (dark brown bars). Shaded areas and errorbars denote SEM. *: p<0.05; **: p<0.01; ***: p<0.001; Wilcoxon ranksum test.

**Figure S5:**
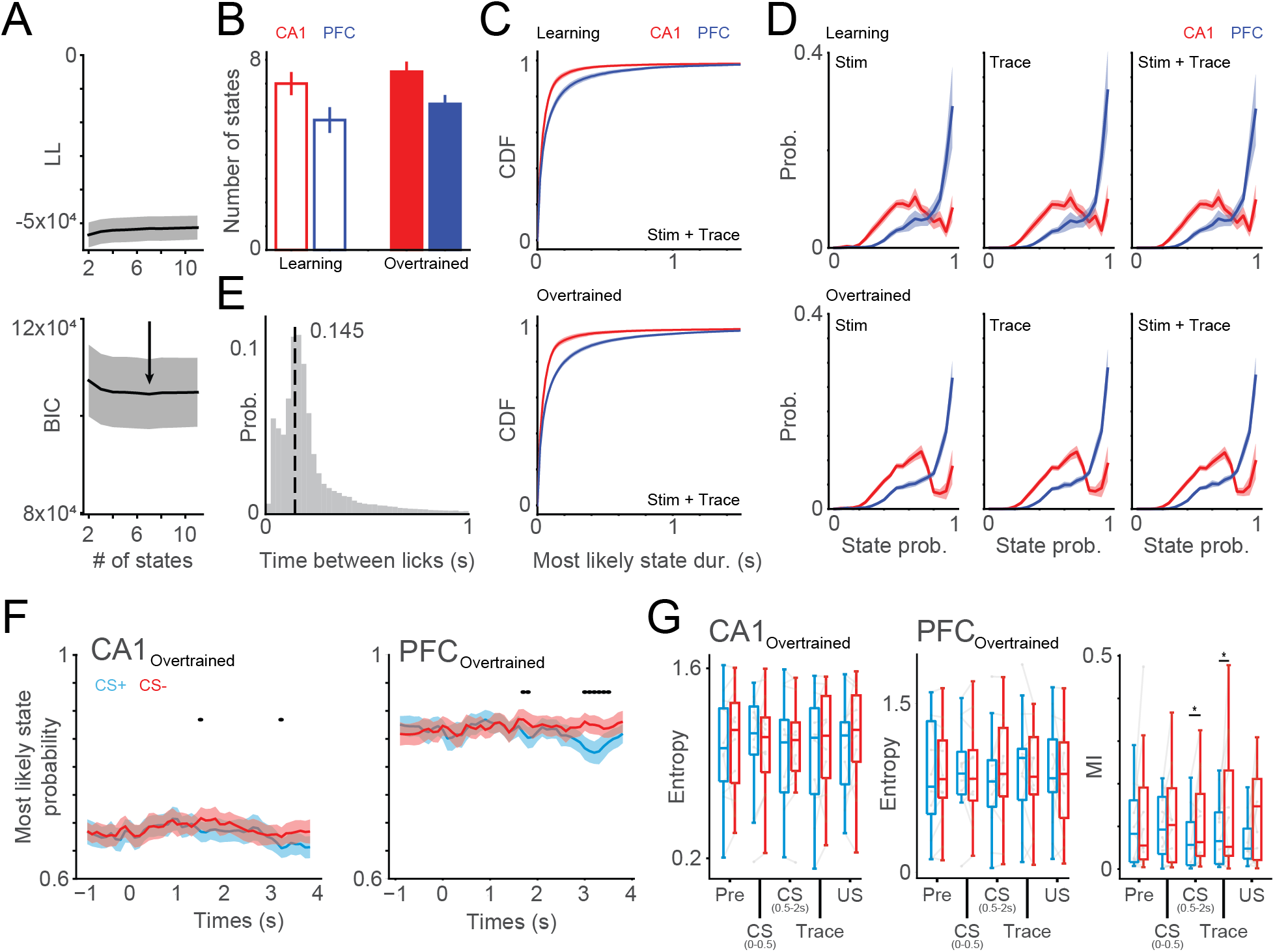
HMM states characterization. **A.** HMM model selection was done using the mean Bayesian information criterion (BIC) over the different cross-validation folds. The model with lowest BIC was used. **B.** Number of states in CA1 and PFC before and after learning. **C.** Cumulative distribution of most likely state duration during CS presentation and trace period. **D.** Most likely state probability distribution during CS, trace and both periods together. Note the higher values in PFC states. **E.** Distribution of the time between consecutive licks. Compare with PFC coding state activation before the first lick. **F.** Most likely state probability over the trial for CA1 (top) and PFC (bottom) overtrained sessions during CS+ and CS− trials. **G.** Entropy (left, middle) and mutual information (right) of most likely state for CA1 and PFC during CS+ and CS− trials.

**Figure S6:**
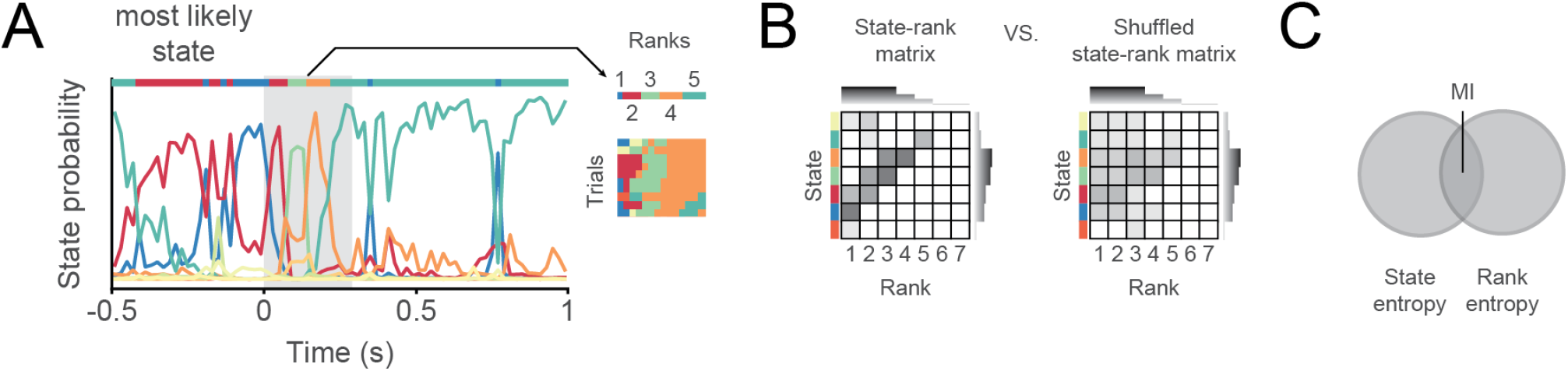
Quantifying state sequences. **A.** Example of state probability along a trial. The most likely state of a window of interest was extracted for each trial, and the rank of states was computed using the order of mean activation time of each state. **B.** State-rank matrix computed using the trials shown in A (left) and an example of a shuffled matrix (right). State-rank matrices indicate how many times a state appear in each rank. Shuffled matrices were computed by randomizing ranks on each trial independently, in order to keep the same number of activation for each state and maximal rank per trial. **C.** The mutual information extracted from the state-rank matrices was an indirect metric of how much information state activation unveils on state rank.

**Figure S7:**
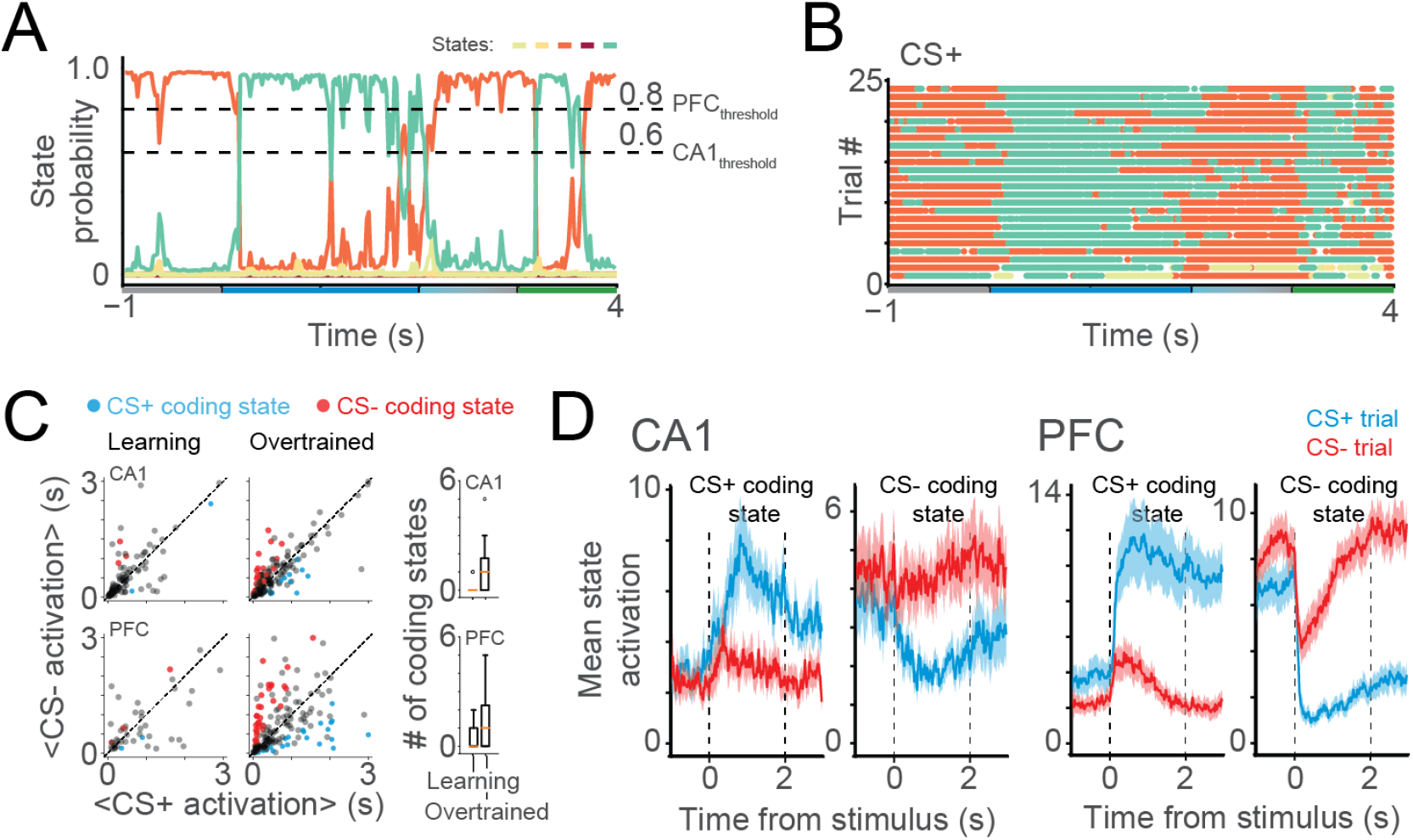
Emergence of CS coding states using conservative thresholds. **A.** Probability of state activation of an example PFC trial. Dashed line show the threshold for CA1 (0.6) and PFC (0.8) used to consider state activation. **B.** Example plot showing activation of states in A over different CS+ trials. **C.** Average CS+ and CS− activation during trace and stimulus period for every state in learning (left) and overtrained (middle) sessions, together with boxplots showing the distribution of coding states per session (right). Coding states appear after learning even when using conservative thresholds. Color denotes CS+ (blue) and CS− (red) coding states. **D.** Average of CS+ and CS− coding states activation over time in both areas. Threshold-defined coding states show a similar profile to the ones defined by the most-likely states.

**Figure S8:**
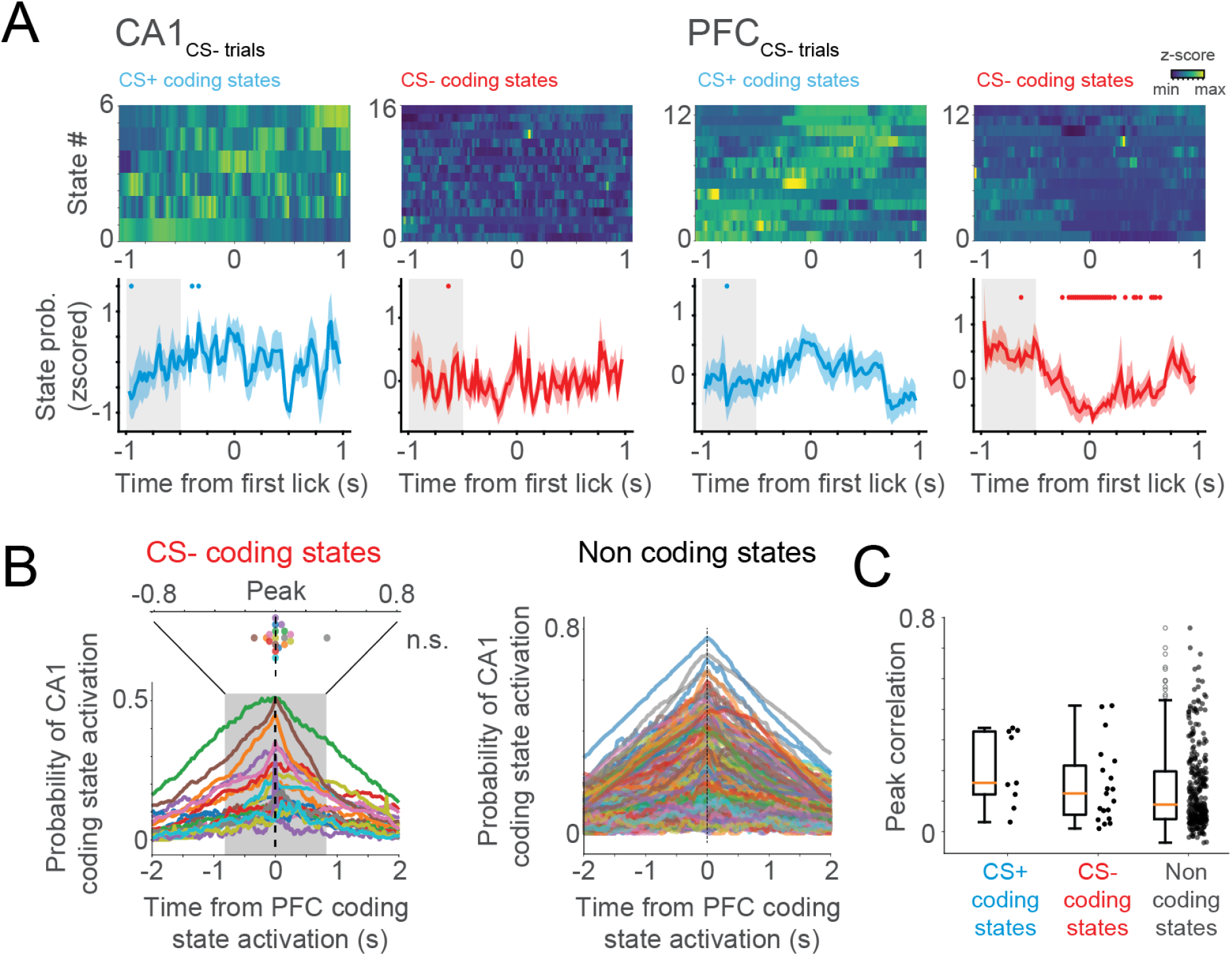
CS coding states in wrong CS− trials and CA1-PFC state correlation. **A.** Coding state’s activation during overtrained sessions and centered on the first lick of the trials. Similar to Figure 7E, but for CS− (wrong) trials. State activation is shown for both CS+ and CS− coding states of CA1 (left) and PFC (right). Notice that PFC coding states seem to predict lick onset even in CS− trials, although this was significant only for the inhibition of CS− coding state. **B.** Cross-correlation between CA1 and PFC CS− and non-coding states activation. Differently from CS+ coding states, CS− and non-coding states in PFC don’t seem to precede CA1 states. **C.** Distribution of the peak correlation values of traces in B. There was no significant difference in correlation between CS+, CS− and non-coding states (Wilcoxon ranksum test).

